# Molecular response to multiple trace element contamination of the European sardine

**DOI:** 10.1101/2024.02.16.580673

**Authors:** Anaïs Beauvieux, Jean-Marc Fromentin, Claire Saraux, Diego Romero, Nathan Couffin, Adrien Brown, Luisa Metral, Fabrice Bertile, Quentin Schull

## Abstract

In marine ecosystems, the presence of trace elements resulting from anthropogenic activities has raised concerns regarding their potential effects on marine organisms. This study delves into the intricate relationship between trace element contamination and the physiological responses of a key marine species in the Mediterranean Sea: the European sardine. Since 2008, this species has been experiencing a significant crisis in the region, prompting numerous studies to investigate the potential factors behind the dramatic decline in sardines’ size, age, and body condition. However, thorough information on chemical contamination by trace elements and its physiological impact on this species was lacking. We found evidence for the accumulation of multiple elements in sardines, with a light East-West contamination gradient within the Gulf of Lions. While macro-physiological parameters (i.e. body condition) were not affected by contamination, pathways involved in cellular organization and response to stress were clearly upregulated, particularly in the liver, but also in muscle. In addition, a global upregulation in processes linked to the immune system, lipid homeostasis and oxidative stress was recorded in the liver. The associated energetic cost may add a substantial burden to sardines that already face multi-factorial constraints. This study also allows to pinpoint biomarkers of exposure and effects that may be important for monitoring Mediterranean sardine’s health. The results of this study and particularly the complex changes in protein expression demonstrate the need for future studies to test the concomitant effects of multiple stressors acting simultaneously, including large scale contamination.

## Introduction

The Mediterranean Sea suffers from high levels of pollution, making it one of the most polluted seas in the world (Gabrielides 1995; Danovaro 2003; Sharma et al., 2021; Robledo Ardila et al., 2024). The sea’s semi-enclosed nature contributes to the accumulation of pollutants, exacerbated by significant atmospheric inputs such as Saharan dust events, intense human activities (Bucchia et al., 2015; Pedrotti et al., 2016) and riverine outflows on the continental shelves (Durrieu de Madron et al., 2011). This is also intimately linked to the enhanced ability of pelagic food chains to bioaccumulate chemical elements in oligotrophic seas such as the Mediterranean Sea (Harmelin-Vivien et al., 2009; Chouvelon et al., 2018). Among natural and anthropogenic pollutants, trace elements are present in all environmental compartments. Some metals (e.g., copper, iron, zinc) are essential for maintaining the healthy function of the cells of all living organisms, particularly due to their role as components or co-factors of different enzymes (Wood et al., 2011). Nonetheless, such elements can become toxic when they exceed certain concentration thresholds, leading to important detrimental effects on the health of organisms (Plum et al., 2010; Dydak et al., 2011; Asaduzzaman et al., 2017). In addition, non-essential trace elements, such as lead, mercury, cadmium, arsenic, nickel, can cause several types of pathological damage at very low concentration (Friberg et al., 1979; Tchounwou et al., 2012; Jaishankar et al., 2014). The accumulation of these chemicals can influence enzymatic and metabolic activities, disrupting ion homeostasis, reducing growth and adversely affecting body condition and swimming performances (Hollis et al., 1999; Bervoets and Blust 2003; Bervoets et al., 2005). Metals are in fact inducers of oxidative stress in many animals (including fish), causing imbalances between the production of reactive oxygen species (ROS) (e.g. H_2_O_2_, HO., O_2_ ^-^, R., ROO) and cell antioxidant activity (Halliwell & Gutteridge, 2015). Increased ROS due to overproduction and/or the inability to destroy them may damage DNA structures and therefore alter the expression patterns of important proteins, hormones, enzymes, etc. (Morcillo et al., 2016; Javed et al., 2017). Metal contamination also modifies the expression of proteins (biomarkers) linked to cellular responses to stress and cell detoxification. The use of biomarkers, such as antioxidant enzymes (Pereira et al., 2013; Williams and Yoshida-Honmachi 2013), heat shock proteins (HSPs, Downs et al., 2006) and metallothioneins (Sakuragui et al., 2013; Williams and Yoshida-Honmachi 2013) to monitor metal pollution has been widely accepted in environmental and ecotoxicology programs.

While experimental data under controlled conditions taught us a lot on the molecular and physiological costs of single contaminants (Isani et al., 2009; Souid et al., 2015; Wang et al., 2013), the challenge that ecologists now face is to understand the effects of trace element mixtures. Indeed, wildlife species are continuously and increasingly exposed to a large number of different chemicals at the same time (Heys et al., 2016). Therefore, the ecotoxicological impacts of trace element mixtures are relatively less understood. Even at low concentrations, trace elements mixed in the environment can have a combined toxicological effect (Heys et al., 2016) forming “reactive chemical cocktails” that explain the synergistic effects of the combination of distinct elements (Kaushal et al., 2018). Considering the effects of only one pollutant at a time can therefore lead to misinterpretation of biomarker data (Celander, 2011).

The emergence of omics tools has provided a powerful means of analyzing complex and integrated responses to contaminants. These tools offer high sensitivity and excellent specificity in studying the molecular changes occurring in organisms (Denslow et al., 2005; Benninghoff 2007). Additionally, high-throughout shotgun proteomics (i.e. the direct and rapid analysis of the entire protein compartment within a complex mixture) provides a unique opportunity to comprehensively examine the expression of thousands of proteins in a specific tissue in a single experiment, using mass spectrometry and bioinformatics techniques instead of traditional biochemical methods (Sanchez et al., 2011; López-Pedrouso et al., 2020). This advance considerably improves our understanding at the protein level and further paves the way for identifying new biomarkers of exposure and effect, which can then be used to develop enhanced monitoring programs to better assess the impacts of pollutants on marine species (Apraiz et al., 2006; Benninghoff 2007).

We applied a quantitative proteomics approach to the European sardine (*Sardina pilchardus*) in the northwestern Mediterranean Sea. This small pelagic fish population has indeed shown a drastic decrease in size and body condition since 2008 (Van Beveren et al., 2014), resulting in a lower value on the stalls, which in turn has led to the collapse of the French small pelagic fish fisheries, which landed less than 200 tons in 2022 against 15,000 tons in the early 2000s (Van Beveren et al., 2016). The hypotheses of top-down control through overfishing or natural predation or epizootic diseases have been refuted (Van Beveren et al., 2016b, 2017, Queiros et al., 2018). The main hypothesis to explain the observed changes is a modification of the quality and/or quantity of sardine preys (such as copepods, Brosset et al., 2016) due to multifactorial environmental changes (SST, Upwelling, Stratification, Convection, WeMO, Chla concentration etc. Feuilloley et al., 2020), thus limiting fish energy resources (bottom-up control, Brosset et al., 2016; Saraux et al., 2019). However, little is known about a potential impact of contaminants on Mediterranean sardines, since few ecotoxicological studies have been performed so far on small pelagic fishes in the Gulf of Lions (but see COSTAS project; Tronczynski et al., 2013 and SUCHIMED campaign; Bouchoucha 2021). In the Mediterranean Sea, surface-enriched concentrations of multiple metals (Co, Cu, Ni and Zn), along with their significant negative correlation with salinity, suggest that concentrations are influenced by the Rhône River plume (e.g. Cossa et al., 2017; Radakovitch et al., 2008). Without a significant tide, the surface extension and main direction of this plume depend on wind and outflow forcing conditions (Broche et al., 1998). In addition, given the sardines’ considerable swimming ability, it is uncertain how the spatial distribution of pollutants can overlap that of sardines. Hence, it is crucial to obtain a thorough knowledge of the contamination pressures exerted on small pelagic species such as sardines, as already highlighted by the United Nations Environment Programme, UNEP (Baker et al., 2013).

The present study therefore focused on assessing in-situ trace element contamination in juvenile European sardines in the Gulf of Lions (NW Mediterranean Sea) and on the impact of contaminant cocktails on individual health using a proteomic approach. The specific objectives of this work were to: (i) assess the trace element load in wild juvenile sardines and evaluate the spatial variability of this contamination; (ii) investigate the impact of trace element mixtures on the proteome of liver (the main organ implicated in xenobiotic metabolism as well as blood glucose regulation, protein synthesis, bile production, vitamin and mineral storage, steroid metabolism, and immune function) and red muscle (an organ with high metabolic capacity contributing to a major part of total muscle respiration rate, see Teulier et al., 2019), and identify potential new biomarkers for both tissues that could serve as valuable tools in the long-term monitoring of European sardines’ health.

## Methods

### Study sites and sampling

We collected 105 juvenile sardines in the Gulf of Lions (northwestern Mediterranean Sea, Figure 1). Specimens were collected in June–July 2021 during the yearly PELMED pelagic survey conducted by the French Institute for the Exploitation of the Sea (Bourdeix & Hattab, 1985). Trawl locations were dependent on the acoustic echoes detected, but covered a wide west-east gradient. Between 3 to 12 sardines per trawl were selected. Sagittal otolith pairs were removed from their otic cavity and read to estimate age. Only age 0 individuals were selected for further analyses. All individuals were measured (total length TL, in mm) usig a metal ruler (±0.1 mm) and weighed (total mass M, in g) using a digital balance (±1 g). Their body condition was calculated using the Le Cren index Kn as estimated by Brosset et al. (2015), which is commonly used as an indicator of general well-being:

**Figure 1.**
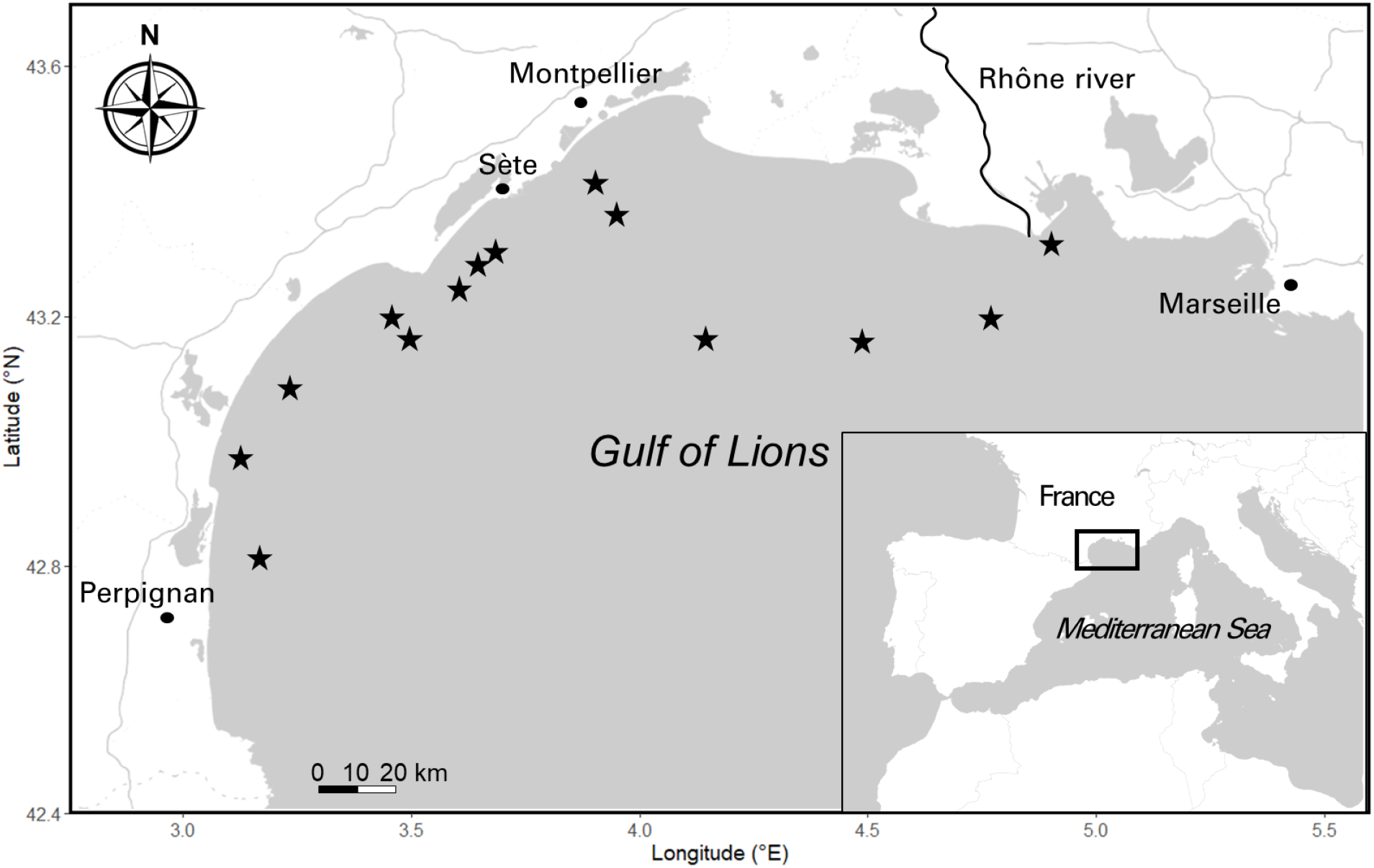
Study area in the Gulf of Lions (northwestern Mediterranean Sea). Pelagic trawling locations for the 105 *S. pilchardus* juveniles in the Gulf are indicated by (⋆).

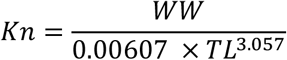

where WW is the total wet mass in g and TL the total length in cm.

Liver and a piece of red muscle were carefully removed to avoid any protein contamination (gloves, a hairnet and a mask were worn by the experimenter and ceramic tools washed with ethanol 96° and MiliQ water), immediately frozen in a dry shipper containing liquid nitrogen (∼-190 °C, ©CX100 Worthington Industries) on-board and later stored at -80 °C at the lab until proteomics analyses. The white muscle on both sides of the fish was removed using cleansed ceramic tools to avoid any metal contamination and stored at -40 °C on-board for the week at sea and then stored at -80° C at the lab until trace element analyses (one side) and lipid content (other side). The red muscle was cleanly separated from the white muscle by scraping the lateral skin stripped from the animal. White muscle samples were taken from the epaxial mass.

### Body condition and reserve lipids

Fish body condition was further investigated by determining white muscle reserve lipid content. Lipids were extracted using roughly 0.1 g of muscle in a solvent mixture of chloroform-methanol 2 : 1, v/v according to the method of Folch et al., (1957) and the content of each class of lipids was measured by chromatography, using a Iatroscan as detailed in Sardenne et al., (2019). Triacylglycerols (TAG), diacylglycerols (DAG as TAG precursors) and free fatty acids (FFA) were grouped together as reserve lipids (Zhol et al., 1995; Tocher 2003; Lloret et al., 2013). The proportions of FFA were also checked to ensure that the lipids had not been degraded during sample storage. Here, the proportions were 0.02% on average (only one sample exhibited a concentration in FFA, whereas all the others showed no presence of FFA), significantly below the 25% limit recommended by Parrish (1988).

### Trace element analysis

All white muscle samples were lyophilized using a freeze dryer (®Leybold, Cologne, Germany). For aluminium (Al), arsenic (As), beryllium (Be), bismuth (Bi), cadmium (Cd), chromium (Cr), copper (Cu), lithium (Li), nickel (Ni), lead (Pb), rubidium (Rb), antimony (Sb), strontium (Sr), titanium (Ti), thallium (Tl) and zinc (Zn) determination, the samples underwent a pre-treatment process where 0.5 grams of each muscle sample were subjected to acid digestion using 4 mL of HNO_3_ (69%) and 1 mL of H_2_O_2_ (33%) in special Teflon reaction tubes within a microwave digestion system (UltraClave-Microwave Milestone®) for 20 minutes at 220 °C, and then diluted to 10 mL with double deionized water (Milli-Q). Samples were analyzed using inductively coupled plasma optical emission spectrometry (ICP-OES, ICAP 6500 Duo, Thermo) to determine the levels of these elements. The detection limit was 0.001 mg/kg. Each sample was read in duplicate and averaged. Based on UNE-EN ISO reference 11885, multi-element calibration standards (SCP Science, in 4% nitric acid) were assembled with different concentrations of inorganic elements. For calibration, see Romero et al. (2020).

Total mercury (Hg) content was measured using an atomic absorption spectrometer AMA254 Advanced Mercury Analyzer (Leco) without pre-treating or pre-concentrating the samples (wavelength = 253.65 nm, detection limit (DL) = 0.003 μg/g). The recovery rate for reference materials (Mercury ICP Standard 1000 mg/L Hg, Merck) was above 95%.

### Quantitative proteomic analysis

In the following proteomic approach, we opted to analyze a subset of 29 individuals who exhibited the most distinct contamination profiles. This selection process involved assigning ranks to sardines based on the level of each contaminant detected. The ranking ranged from 1 (indicating the lowest contamination level) to 105 (representing the highest contamination level). The individuals were then selected based on the total sum of these ranks. Consequently, we identified two groups, one comprising 14 sardines with the lowest contamination levels and the other consisting of 15 sardines with the highest contamination levels (see Statistical analysis section for further details on individuals’selection). Proteomic analyses of liver and red muscle were conducted as two separate experiments. The process of sample preparation, nanoLC-MS/MS analysis, and mass spectrometry data analysis is described in detail in ESM1. The procedure involved protein extraction and electrophoresis using SDS-PAGE, followed by in-gel digestion with trypsin (Promega, Madison, WI, USA). The resulting peptides were extracted and analyzed using a nanoUPLC system (nano-Acquity, Waters, Milford, MA, USA) coupled to a quadrupole-Orbitrap hybrid mass spectrometer (Q-Exactive HF-X, Thermo Scientific, San Jose, CA, USA), controlled by XCalibur software (v4.0.27.9; Thermo Fisher Scientific). The Q-Exactive HF-X was operated using a data-dependent acquisition (DDA) strategy by selecting the Top-20 most intense ions in MS1 for fragmentation in MS2. MS raw data processing was performed in MaxQuant (v2.0.3.1) (Cox et al., 2014), using Andromeda algorithm to search peaklists against a protein database containing protein sequences from a home-made annotation of the genome of the European sardine (TaxID: 27697; Genbank assembly accession: GCA_003604335.1; Assembly Name: UP_Spi). Only the proteins identified with at least two peptides were retained. Protein quantification was performed using unique peptides only via the MaxLFQ option implemented in MaxQuant (Cox et al., 2014) which is a label-free quantification (LFQ) method that determines protein abundance based on the intensity of peptide signals. The mass spectrometry proteomics data have been deposited to the ProteomeXchange Consortium via the PRIDE (Perez-Riverol et al., 2019) partner repository with the dataset identifiers PXD037276 (liver) and PXD037313 (red muscle). QC-related measurements indicated stable performances of the analysis system all along the two experiments, with median coefficients of variation (CV) of 0.47% (liver) and 0.56% (red muscle) for retention times of iRT peptides over all injections. Median CVs of only 10.9% (liver) and 13.8% (red muscle) were obtained for LFQ values of the proteins quantified from the repeated injections of reference samples.

### Protein functional annotation

To analyse our proteomic data from a functional point of view, we examined the functional annotations of proteins listed by the AMIGO consortium (Gene Ontology, GO; http://geneontology.org/). However, only one sardine protein is annotated in GO databases. We therefore took advantage of the crucial evolutionary position of the spotted gar (*Lepisosteus oculatus*) between teleost fishes and humans, which creates a very valuable bridge between their genomes (Braasch et al., 2016), and of the fact that gar proteins are well annotated in GO databases. We began by searching *L. oculatus* sequences (UniprotKB, 22,463 protein entries; September 2022) for proteins homologous to *S. pilchardus* proteins using BLAST searches (FASTA v36.1.4 program; downloaded from http://fasta.bioch.virginia.edu/fasta_www2/fasta_down.shtml), and only the top BLAST hit for each protein was retained. From *L. oculatus* protein sequences, the same strategy was employed to search for human homologous proteins (TaxID: 9606; Reference proteome accesed in September 2022). The relevance of the match between homologous proteins was checked manually and only 57 of 2505 and 22 of 1112 matches (liver and red muscle, respectively) could not be validated. An automatic extraction of GO annotations was then performed using the MSDA software (Carapito et al., 2014).

### Statistical analyses

All statistical analyses were performed with the statistical open source R software v.4.1.1 (R Core Team, 2021). Element concentrations below Limit Of Detection (LOD, 0.001 mg/kg) were set to “LOD/ √2” as suggested by Verbovsek (2011) as the best substitution method.

Analyses comparing inorganic contamination patterns included only metals that were detected in at least 50% of individuals out of the 105 collected. In order to consider possible relationships between contamination and fish body size, we used a linear model (LM) between each contaminant concentration and the individual total length. Normality and homoscedasticity of the residuals were verified using the Shapiro’s and Levene’s tests, respectively. When distribution of the residuals did not follow a normal distribution, a log transformation was applied prior to model fitting. If necessary (P < 0.05), size-corrected contaminant concentration was defined as back-transformed residuals of the regression model.

To investigate relationships between contaminants and to evaluate a possible influence of the Rhône River on trace element loads in sardines within the Gulf of Lions, we performed a Principal Component Analysis (PCA) on scaled size-corrected contaminant concentrations of the 105 individuals and projected distance from the Rhône River mouth on the PCA. We also assessed Spearman correlation between the three first principal components (PCs) and the distance of the individuals from the Rhône River delta. To evaluate a potential effect of inorganic contamination on individual’s condition, we also performed LMs between the first three PCs and body condition and reserve lipids.

A second PCA approach was performed using only the 29 individuals selected for proteomic analysis as objects, while the descriptors were scaled size-corrected contaminant concentrations. The aim of this analysis was to summarize all the information to investigate the relationships between contaminants and individual proteomes. PCA axes (PC1, PC2, PC3) were retained as principal components reflecting contaminant mixture 1, mixture 2, mixture 3 and used in further analyses.

Prior to statistical analysis, the proteomic datasets (liver and red muscle) were filtered to keep only proteins that were expressed in at least 70% of individuals in each contamination group (common threshold used in existing literature (Quque et al., 2023; Wang et al., 2020) leaving 2115 (out of 2505) and 913 (out of 1112) expressed proteins in the liver and red muscle, respectively. Three and one individuals were removed due to a high number of missing values for liver and muscle proteome analysis, respectively (supported by the missing value heatmap and proteomic clustering isolation of these individuals). Remaining missing values were imputed for each contamination group based on random forest method (Jin et al., 2021; Kokla et al., 2019). Previous studies have shown that this method exhibits strong performance and is particularly well-suited for label-free proteomic studies, where the underlying reasons for missing data are not fully understood (Jin et al., 2021; Kokla et al., 2019). A log 2 transformation was used to normalize the protein intensities.

### Co-expression network and enrichment analysis

#### Gene co-expression network and hubproteins identification

Weighted gene co-expression network analysis implemented in the WGCNA R package (Langfelder & Horvath, 2008) was used for both tissues (liver and red muscle) to identify groups, or modules, of proteins whose expression significantly correlated with inorganic contamination in white muscle. First, an adjacency matrix for all pairs of proteins was constructed using the Spearman correlation raised to the power (beta) of seven and six to approximate a scale-free network for liver and red muscle, respectively (Langfelder & Horvath, 2008). The adjacency matrix was then transformed into a topological overlap dissimilarity matrix and a combination of hierarchical clustering and a dynamic tree-cutting algorithm were used to first define and then merge co-expressed modules of proteins. Proteins outside any module (indicating low co-expression) were gathered in a grey module. The module’s eigengene (i.e. first module eigenvector: first axis of a principal component analysis conducted on the expression of all module proteins) summarizes the expression of all proteins in that module. Then, we investigated whether the eigengene of the modules was correlated with each contaminant mixture (identified through the PCA analysis, see above), using Spearman correlation. Modules with a significant (P < 0.05) correlation and at least R ≥ 0.4 were retained for further analysis (only the grey module, indicating low co-expression, for muscle presented a significant correlation below 0.4). Positive and negative relationships reflect the up- and down-regulation of the module with an increasing concentration of a contaminant mixture, respectively. To explore the functional and physiological mechanisms associated with each module, we then performed functional enrichment analyses.

#### Functional enrichment analysis and pathway network

For both tissues, enrichment analysis (biological process and cellular component) of the proteins constituting each module was performed using Fisher’s exact test using the ‘*GO_MWU’* package particularly suitable for non-model organisms (https://github.com/z0on/GO_MWU) (Dixon et al., 2015). The background used for each tissue consisted of the set of proteins identified in that specific tissue, ensuring two distinct backgrounds for the analysis. This approach prevents biased enrichment from a generic background, ensuring accurate determination of the functional significance of the identified modules (Wijesooriya et al., 2022). Enriched GO terms are displayed in a dendrogram plot with distances between terms reflecting the number of shared proteins. Additionally, we assessed the Spearman correlation between each protein and the contaminant mixtures (PCA axes) to infer “gene significance” (GS). Focusing on the top 10 enriched biological process terms (GO terms), these GS relationships were compiled within a circle plot for each enriched pathway of the module. These plots highlight the up-or down-regulation of the proteins and pathways in response to the contaminant mixtures.

#### Network visualization

ClueGO was used to illustrate overrepresented Gene Ontology terms. ClueGO is a Cytoscape plug-in that visualizes the non-redundant biological terms for large numbers of proteins and integrates the GO terms to create a GO/pathway network (Bindea et al., 2009). A network for each tissue was made in order to visualize the pathways impacted by the three contaminant mixtures.

#### Hubproteins identification

For each module, GS was correlated with module membership (MM; Spearman correlation of protein intensity versus the module eigengene) to identify proteins showing the highest degree of connectivity within a module and with the considered contaminant mixture (hubproteins). The top 5 proteins with simultaneously the highest MM and GS (rank sum) were selected as hubproteins in each module. Due to their central position in the network, hubproteins are expected to play important biological roles within their module and are considered as potential biomarkers.

#### Functional comparison between tissues

For each contamination mixture, the similarity of the functional response between tissues was compared by plotting their respective GO term delta ranks against each other. GO term delta ranks correspond to the difference between the mean ranks based on the GS of the proteins the GO term contains *vs* the mean ranks of all the protein not included. Positive and negative delta ranks indicate that the GO term tends to be regulated upwards or downwards, respectively. For a given pathway, the strength of the relationship reflects the similarity of the enrichment between liver and red muscle. It is important to note that these plots do not represent a formal statistical test, as the data points (gene ontology categories) are not independent. Indeed, they often encompass overlapping sets of proteins, nonetheless it allows identifying functional similarity or dissimilarity between functional enrichments.

## Results

### Trace element burden across Gulf of Lions

Thirteen trace elements (Al, As, Cu, Cr, Hg, Li, Ni, Pb, Rb, Sr, Ti, Tl and Zn, ESM2) were retained after selection of contaminants detected (> LOD) in at least 50% of individuals (i.e. 53 individuals). Only three elements (Pb, Rb and Tl) exhibited a significant relationship with fish length and were size-corrected. While Tl displayed a negative relationship with length (R^2^ = 0.16, P < 0.001), Pb and Rb had a positive association (R^2^ = 0.10, P = 0.009 and R^2^ = 0.09, P = 0.001 respectively).

Considering the 105 sardines sampled across the Gulf of Lions within the same PCA, the first three principal components explained 24.2%, 15.3% and 12.8% of the variance, respectively (around 52% in total, ESM3a and Figure 2). The main contributing variables for the first axis (PC1) were Zn, Sr, Rb, Cu, Pb and As, which all displayed positive contributions (ESM3b and Figure 2). Ni, Cr were positively related to PC2 while, to a lesser extent, Tl was negatively loaded (ESM3c and Figure 2). Ti and Al were negatively loaded along the third axis in opposition to Cr, Ni, Tl and Li (ESM3d and Figure 2). Highlighting the distance of the various sampling sites from the mouth of the Rhône River within the same PCA showed a slight east-west contamination gradient (Figure 2). PC1 and PC2 were negatively correlated with the distance from the Rhône river, albeit slightly above significance levels (ESM4). Nonetheless, there was considerable inter-individual variability in trace element contamination, which probably blurred this east-west gradient (ESM4). The three first PCs displayed no significant correlation with individuals’ body condition (P = 0.47, P = 0.18 and P = 0.57 for PC1, PC2 and PC3, respectively), nor with the reserve lipid content (P = 0.55, P = 0.66 and P = 0.79 for PC1, PC2 and PC3, respectively).

**Figure 2.**
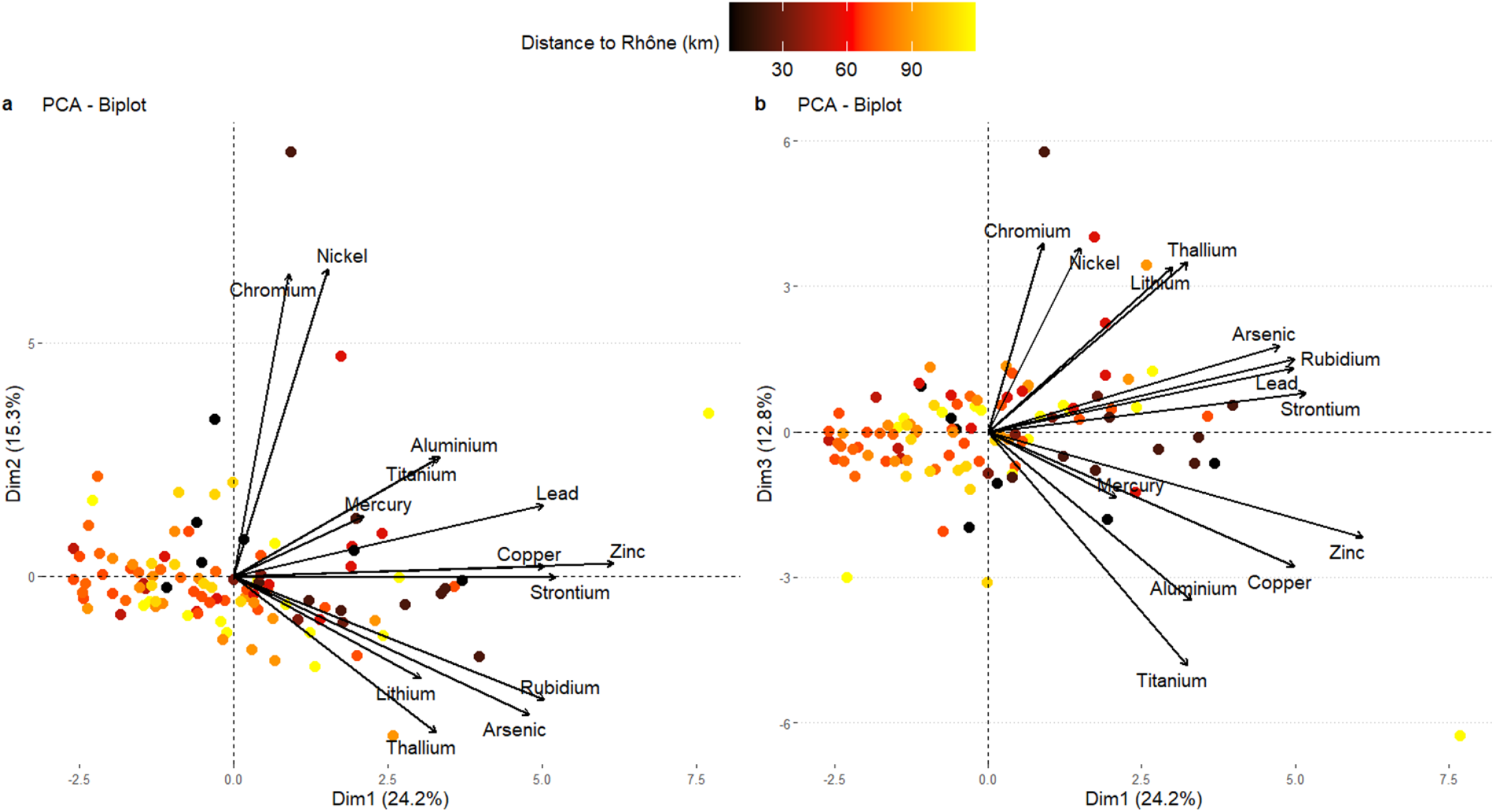
Biplot of PC1 vs PC2 (a) and PC1 vs PC3 (b) of the PCA built with size-corrected level of trace elements present in the 105 individuals sampled across the Gulf of Lionss. Each point represents an individual. Colour gradient indicates the distance from the Rhône river mouth.

Regarding the second PCA performed on the 29 individuals selected for proteomic analysis, the first three principal components explained 33.3%, 16.9% and 13.6% of the total variance, respectively (around 64% in total, Figure 3 and ESM5). All 13 contaminants loaded positively on PC1 with 8 of them contributing strongly. Further, PC1 clearly separated the 2 groups of individuals previously selected (high contamination *vs* low contamination). Therefore, PC1 was used as a proxy for general inorganic contamination level, which allows to work along a contamination gradient instead of categorical contaminations. PC2 was mostly determined by a strong positive contribution of Ni and Cr, while PC3 was determined by a strong positive contribution of Ti and Al and marginally by a negative contribution of Li.

**Figure 3.**
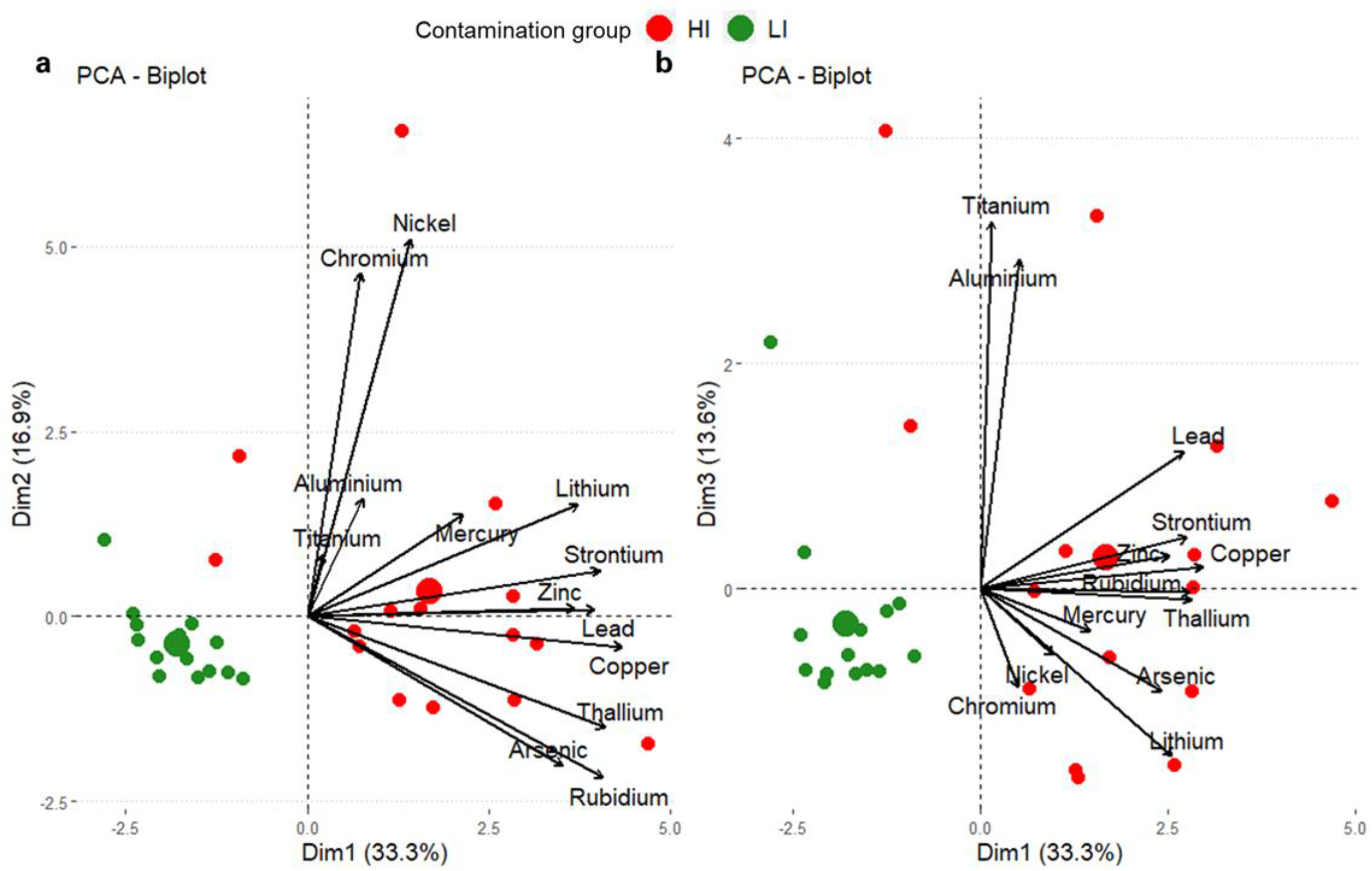
Biplot of PC1 vs PC2 (a) and PC1 vs PC3 (b) of the PCA built with level of inorganic contaminants present in the 29 individuals selected for proteomic analysis. Each point represents an individual. Colours indicate contamination level of individuals (HI: High contamination and LI: Low contamination). The larger circles represent the barycenter of the individuals for a given group.

The three PCs independently identified contaminant mixtures, which are listed in Table 1. We next used proteomics to investigate the potential impact of these three contaminant mixtures on the health of the Mediterranean sardine population.

**Table 1.** Main variables that contribute to the first three principal components of the PCA on the 29 individuals selected for proteomic analysis. Each PC represents a contaminant mixture.

| Mixture 1 - PC1 |  | Mixture 2 - PC2 |  | Mixture 3 - PC3 |  |
| --- | --- | --- | --- | --- | --- |
| - | + | - | + | - | + |
| none | Cu, Tl, Rb, Sr,<br>Pb, Li, Zn, As | none | Ni, Cr | Li | Ti, Al |

### Physiological response to inorganic contamination

Using a co-expression network analysis, we focused on the impact of metal contamination at the proteome level on two key organs in fish (the liver and red muscle).

#### Liver proteome

We detected 9 distinct modules (WGCNA analysis) containing a total of 2115 proteins. We then investigated the relationship between expression modules and the three metal mixtures to assess the potential health effects of inorganic contaminant exposure. Both mixtures 1 and 3 displayed significant positive correlations (P < 0.05, encompassing approximately 31% of the detected proteins) with the same two co-expressed modules (turquoise and yellow, Figure 4), reflecting a general overexpression of proteins within these 2 modules when mixtures 1 and 3 increased in concentration. Mixture 2 did not display any relationship with any of the co-expressed modules.

**Figure 4.**
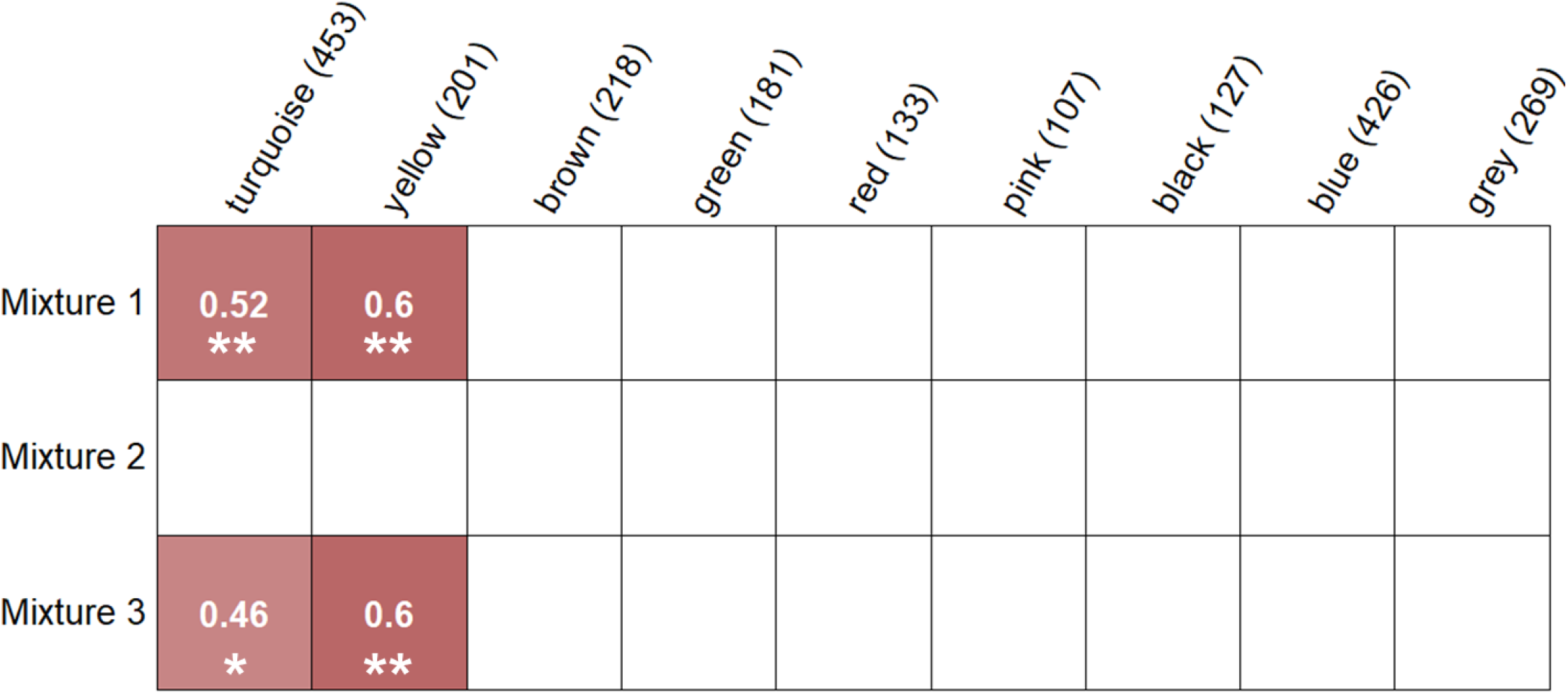
Correlations between module eigengenes (columns) and trace element mixtures (rows) for liver proteome. Values in parentheses with module names indicate the number of proteins belonging to each module. Values in the cells are Spearman’s correlation coefficients. Blank cells indicate non-significant correlation (P>0.05). *: P < 0.05, **: P < 0.01

The positive correlation of the turquoise module with mixture 1 (PC1, R = 0.52, P < 0.01) and mixture 3 (PC3, R = 0.46, P < 0.05) involved an overall upregulation of 170 GO terms (FDR < 0.01), mainly involved in anatomical structure and development (e.g., epithelial cell development (GO:0002064), regulation of anatomical structure size (GO:0090066), response to stimulus/stress (e.g. defense response (GO:0006952), cellular response to DNA damage stimulus (GO:0006974), protein metabolism (e.g. positive regulation of gene expression (GO:0010628), regulation of protein polymerization, transport and cellular organization (e.g. cell development, regulation of cytoskeleton organization and cell cycle process (spindle organization, cell cycle process, Figure 5 and ESM6a). Proteins from the turquoise module were part of multiple subcellular compartments, such as the nuclear periphery, endocytic vesicles, immunological synapses and the cytoskeleton (ESM6b). The positive correlation of the yellow module with mixture 1 (P < 0.001) displayed a significant enrichment in 71 pathways mainly related to the immune system, the metabolism of lipids and lipoproteins and transport (Figure 5 and ESM7a). Most of these pathways were upregulated in response to an increase in mixture 1 (i.e. Cu, Rb, Sr, Tl, Zn, Li, Pb and As). As for the cellular component analysis, proteins were mainly located in lumens (vesicle, endoplasmic, endocytic) and were subunits of protein complexes (protein-lipid complex, hemoglobin complex, ESM7b).

**Figure 5.**
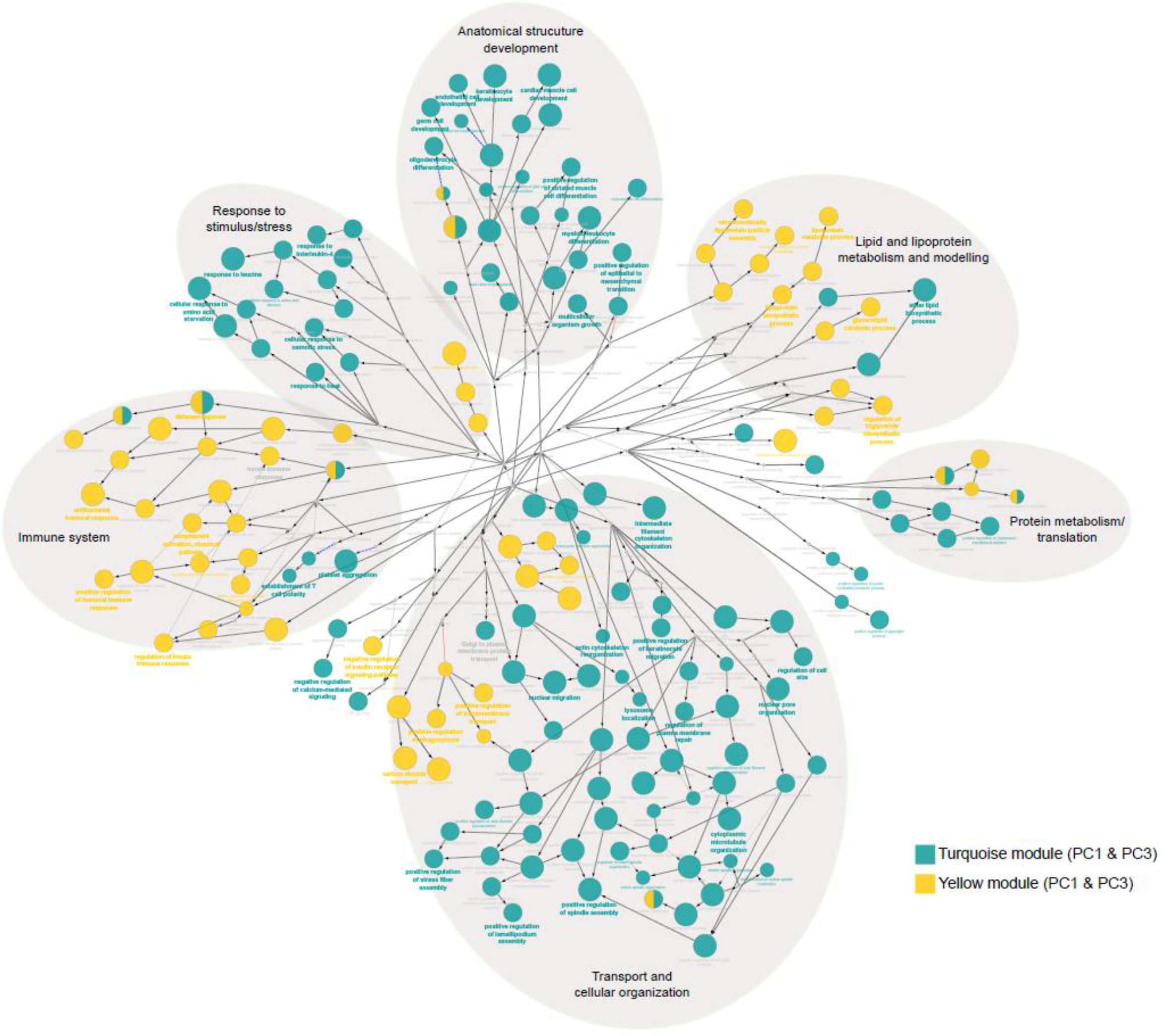
Significant GO terms (FDR < 0.001) and ontological relationships in the liver. Colored circle size reflects the level of FDR-adjusted statistical significance. Terms enriched in the same module were grouped and presented in the same color. Each leading term, which has the highest significance, is indicated by bold font. Biological processes were grouped in larger pathways (large grey circles).

To identify potential biomarkers, hubproteins (top 5 proteins with the highest GS and MM) were selected for each module (see Table 2). For the turquoise module, hubproteins were mainly implicated in protein or lipid transport (Table 2). Interestingly, one of them, the serine/threonine-protein kinase mTOR, was involved in multiple pathways such as cell development, cell structure and metabolism. Potential biomarkers for the yellow module included 2 apolipoproteins with one presenting antioxidant activity (i.e., the Apolipoprotein A-IV). The three other hubproteins were involved in diverse pathways such as protein synthesis with the eukaryotic translation initiation factor 2 subunit 3, cytoskeleton and protein transport with the spectrin beta chain and the DnaJ homolog subfamily C member 13 which is not only implicated in protein transport but also exhibits chaperon function.

**Table 2.** Top 5 hubproteins selected for each co-expression module in the liver proteome. The name and UniprotKB ID are indicated for each of them as well as their main functions, their module, their expression pattern (up or down-regulation), contaminant mixture they correlate with and the module membership.

| Name | UniprotKB ID | Module | Main function | Expression pattern | Principal Component | Module membership |
| --- | --- | --- | --- | --- | --- | --- |
| Apolipoprotein B-100 isoform 3 | APOB | turquoise | Lipid transport and metabolism | ↑ up | PC1 and PC3 | 0.83 |
| Serine/threonine-protein kinase mTOR | MTOR | turquoise | Cell growth, Cell cycle progression, Metabolism, Cellular structure | ↑ up | PC1 and PC3 | 0.81 |
| Cytoplasmic dynein 1 heavy chain 1 isoform 5 | DYNC1H1 | turquoise | Intracellular motility: cell cycle and transport | ↑ up | PC1 and PC3 | 0.78 |
| U5 small nuclear ribonucleoprotein 200 kDa helicase | SNRNP200 | turquoise | mRNA processing/splicing | ↑ up | PC1 and PC3 | 0.89 |
| Cytoplasmic dynein 1 heavy chain 1 isoform 2 | DYNC1H1 | turquoise | Intracellular motility: cell cycle and transport | ↑ up | PC1 and PC3 | 0.80 |
| Apolipoprotein A-IV | APOA4 | yellow | Lipid transport and metabolism, Antioxidant capacity | ↑ up | PC1 and PC3 | 0.88 |
| Apolipoprotein B-100 isoform 1 | APOB | yellow | Lipid transport and metabolism | ↑ up | PC1 and PC3 | 0.79 |
| Eukaryotic translation initiation factor 2 subunit 3 | EIF2S3 | yellow | Translation | ↓ down | PC1 and PC3 | -0.77 |
| Spectrin beta chain, non-erythrocytic 1 | SPTBN1 | yellow | Cellular structure, Protein transport/localization | ↑ up | PC1 and PC3 | 0.84 |
| DnaJ homolog subfamily C member 13 | DNAJC13 | yellow | Chaperone, Protein transport | ↑ up | PC1 and PC3 | 0.88 |

#### Muscle proteome

The WGCNA analysis of the red muscle proteome, containing 913 proteins, resulted in four modules namely blue (372 proteins), brown (119 proteins), turquoise (398 proteins), and grey (24 proteins).

Only the blue module, which accounts for 41% of all detected proteins in the red muscle proteome, correlated (positively) with one contaminant mixture (i.e., mixture 1, Figure 6). This module was enriched in cellular organization and ion transport pathways (Figure 7 and ESM8). According to the cellular component enrichment analysis, proteins were mainly located in membranes (ESM8b).

**Figure 6.**
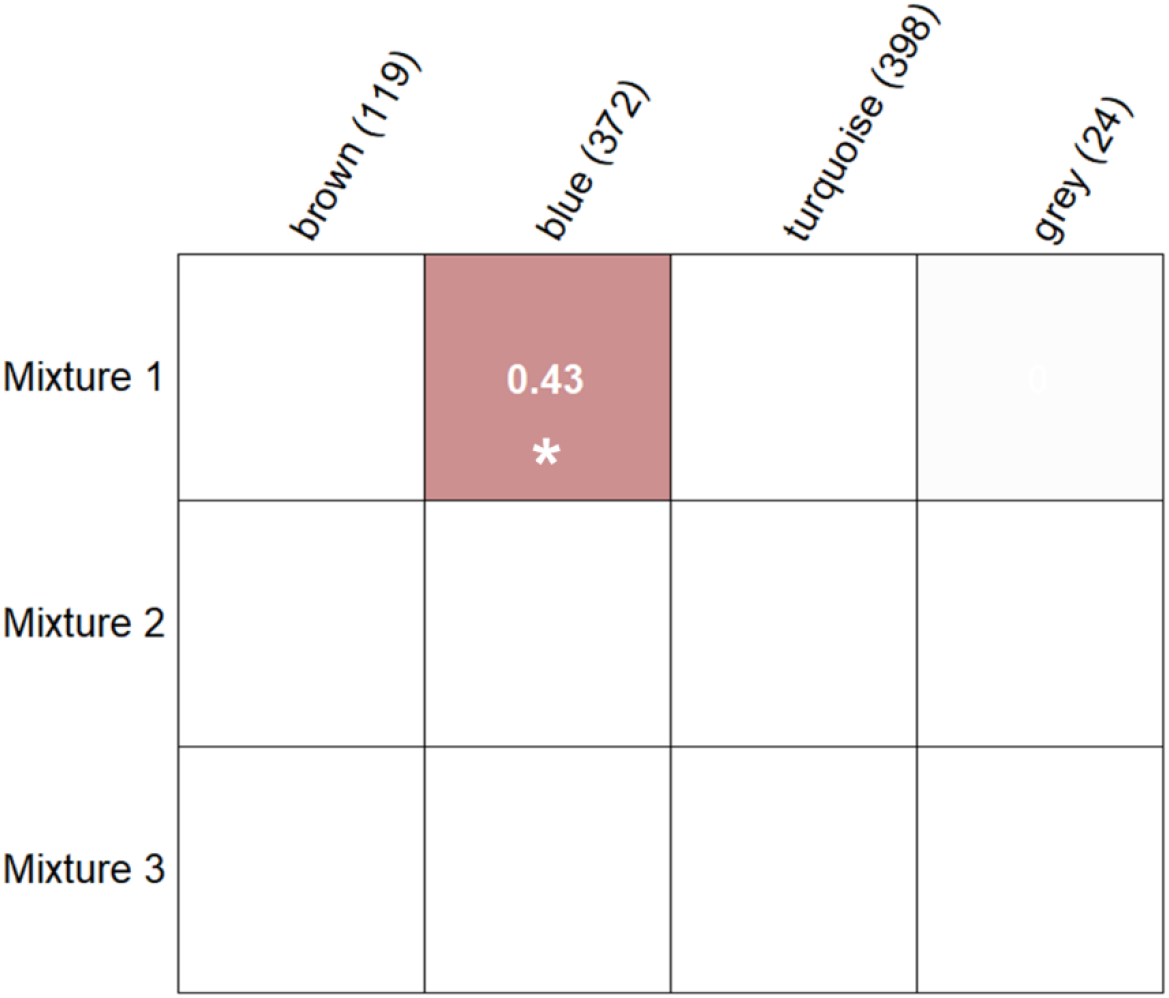
Correlations between module eigengenes (columns) and trace element mixtures (rows) for the red muscle proteome. Values in parentheses with module names indicate the number of proteins belonging to each module. Values in the cells are Spearman’s correlation coefficients. Blank cells indicate non-significant correlation (P > 0.05). *: P < 0.05, **: P < 0.01

**Figure 7.**
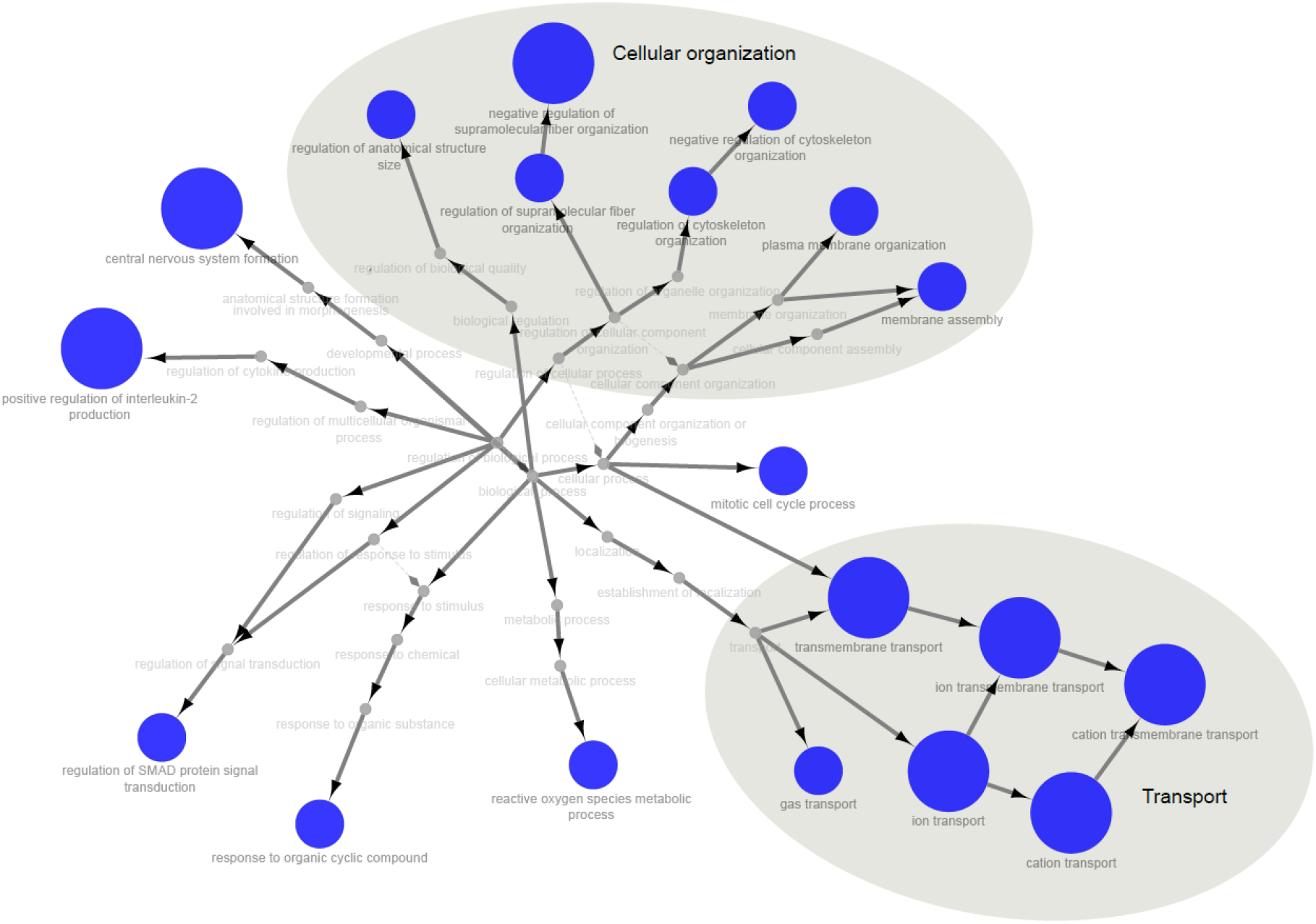
Significant GO terms (FDR < 0.001) and ontological relationships in red muscle. Circle size indicates the level of FDR-adjusted statistical significance. Biological processes were grouped in larger pathways (large grey circles).

Two identified hubproteins for the blue module have a known function in the response to oxidative stress (Gluthatione S-transferase and Hemoglobin subunit zeta), whereas two others are involved in cellular structure and muscle contraction (myosin-binding protein H and Myomesin-2). The last identified hubprotein was the ADP/ATP translocase 1 which is implicated in energy metabolism and apoptosic signaling pathway (Table 3).

**Table 3.** Top 5 hubproteins selected for each co-expression module in the red muscle proteome. The name and UniprotKB ID are indicated for each of them as well as their main functions, their module, their expression pattern (up or down-regulation), contaminant mixture they correlate with and the module membership.

| Name | UniprotKB ID | Module | Main function | Expression pattern | Principal Component | Module membership |
| --- | --- | --- | --- | --- | --- | --- |
| Glutathione S-transferase | LANCL1 | blue | Contaminant detoxification, Oxidative stress metabolism | ↓ down | PC1 | -0.92 |
| Hemoglobin subunit zeta | HBZ | blue | Gas transport, Oxidative stress metabolism | ↑ up | PC1 | 0.90 |
| Myosin-binding protein H | MYBPH | blue | Cell adhesion, Muscle contraction | ↓ down | PC1 | -0.92 |
| ADP/ATP translocase 1 | SLC25A4 | blue | Transmembrane transporter, Apoptotic signalling pathway | ↓ down | PC1 | -0.92 |
| Myomesin-2 | MYOM2 | blue | Cellular organization, Muscle contraction | ↓ down | PC1 | -0.91 |

#### Comparison of liver vs red muscle proteome response

Comparison of the delta-rank values obtained from GO enrichment analysis for both tissues against mixture 1 revealed a significant positive correlation (R = 0.22, P < 0.001 ESM9a). Liver and muscle enrichment for mixture 2 displayed no significant correlation (R = 0.02, P = 0.96, ESM9b), while enrichment for mixture 3 revealed a significant but weak positive correlation between the two tissues (R = 0.06, P < 0.01, ESM9c). This highlights that similar processes appear involved in liver and red muscle in response to contaminant mixture 1 and, to a lesser extent, mixture 3.

## Discussion

Multiple studies have investigated the potential drivers of the drastic decrease in size, age and body condition of sardines in the Gulf of Lions since 2008 (e.g. Brosset et al., 2015, 2016, 2017; Le Bourg et al., 2015; Van Beveren et al., 2016, 2017; Feuilloley et al., 2020). However, thorough information on chemical contamination by trace elements and its physiological impact on this species was lacking. We found evidence for the accumulation of multiple elements in sardines, with a light east-west contamination gradient within the Gulf of Lions. While macro-physiological parameters (i.e. body condition) were not affected by contamination, pathways involved in cellular organization and response to stress were clearly upregulated, particularly in the liver, but also in muscle. In addition, a global upregulation in processes linked to the immune system and lipid homeostasis was recorded in the liver. We are proposing biomarkers of the effects of trace element stress on health, which could serve as a starting point in biomonitoring programs.

### Contamination load in the Mediterranean sardine of the Gulf of Lions

The impact of contaminating loads from the Rhône River on the organisms living in the Gulf of Lions has already been reported with regard to radionuclides, such as ^210^Po (Strady et al., 2015), heavy metals (Cd, Pb and Hg for [60–200 μm] size fraction, Chouvelon et al., 2019) and organic contaminants, such as Polychlorinated biphenyls (PCBs) in plankton (Alekseenko et al., 2018). In addition to the Rhône River inputs, pollutant discharges are also likely to come from industrial and urban activities of the nearby cities of Marseille and Fos-sur-Mer. Small pelagic planktivorous fish such as sardines in the eastern areas of the Gulf of Lions might therefore be highly exposed to chemical contamination, notably via trophic pathways (i.e. plankton). Interestingly, although large inter-individual variability in contamination was recorded in our study, we noted a moderate east to west contamination gradient in the Gulf of Lions. These results differ from previous work (Chouvelon et al., 2019) that did not detect a spatial pattern in trace element contamination in the same species and location. These differences might be due to the targeted tissue, as they considered contamination in the whole organism (liver, muscle, gonad and remaining tissues) whereas we focused on muscle contamination. Indeed, factors such as tissue-specific metabolic processes, affinity for contaminants and turnover rates can contribute to distinct contamination patterns observed across differents tissues. Another reason for discrepancies between our and previous studies might involve the fact that individuals were sampled at different life stages in those former studies, whereas we focused on individuals in their first year of life (age 0). Indeed, confounding effects with contaminant elimination through reproduction as well as bioaccumulation with age may explain large variations and no clear spatial pattern (Bodiguel et al., 2009). Canli and Atli (2003) highlighted the importance of fish size in determining the rate of physiological processes, as size may influence the uptake, distribution, and elimination of trace elements. In our study, only three trace elements (out of 13) displayed significant relationships with fish size, namely Pb, Rb and Tl. This lack of relationship with size for most of trace elements could result from the complexity of bioaccumulation processes involving interaction between various routes of absorption, excretion, passive release, and metabolization (Joiris et al., 1999; Guo et al., 2016; Tang et al., 2017). Yet, the study of one-year-old gilthead seabream juveniles (Beauvieux et al., 2024) showed comparable outcomes. This indicates that when examining a single age group, size differences are likely minimal, simplifying the study of spatial contamination. That said, exploring multiple age groups remains essential to uncover intricate bioaccumulation patterns.

Trace metal concentrations in Mediterranean surface waters and top predators are generally higher than in the Atlantic Ocean (Boyle et al., 1985; Cossa and Coquery 2005). This phenomenon can be largely attributed to the Mediterranean’s semi-enclosed geography, which promotes pollutant accumulation. This accumulation is further exacerbated by significant atmospheric inputs, such as Saharan dust events, intense human activities (Bucchia et al., 2015; Pedrotti et al., 2016), and riverine discharges on continental shelves (Durrieu de Madron et al., 2011). Additionally, there is evidence suggesting that the Mediterranean pelagic food webs may have a heightened capacity to bioaccumulate certain trace elements, particularly Hg (Cossa and Coquery 2005; Harmelin-Vivien et al., 2009; Chouvelon et al., 2018). This pattern also turned out to be true for As, Ni, Cr, Pb, Zn and Hg in sardines recorded in numerous studies (considering only the European sardine in ESM10). Trace element concentrations found in the present study were of a similar order of magnitude to those found in other studies focusing on the same or close species of the Mediterranean, Atlantic or Indian Oceans, except for Li. Averaged Li levels were 32 times higher than those found by Guérin et al., (2011) on the Mediterranean sardine (but Li levels were measured in only 4 individuals in this previous work) and 2 times higher than those found by Lozano-Bilbao et al., (2019) on the European sardine in the Atlantic Ocean.

Similarly to findings in seabreams (Beauvieux et al., 2024), we identified a lack of monitoring for several elements including Li, Rb, Sr, Ti and Tl in all locations (Indian and Atlantic Oceans and the Mediterranean Sea, ESM10 and ESM11). This is partially explained by the absence of regulatory limits for these elements in fish muscle (along with Cr and Ni), despite their increasing use in multiple areas (industrial, agriculture, high-tech and medication industries) and despite their toxicity for organisms (Exley et al., 1991; Authman 2011; Aslam and Yousafzai 2017; Blewett and Leonard 2017; Kumari et al., 2017; Genchi et al., 2021).

The European regulation sets contamination thresholds for only three specific inorganic trace elements, Pb, Cd and Hg. In contrast, other nations, such as Turkey and Greece have adopted thresholds for a larger set of contaminants (for a comprehensive review, refer to Guerra-García et al., 2023). Notably, trace element levels in sardine specimens from the present study remained below the threshold values for fish consumption set by most international regulations except for As that exceeded these limits. These findings emphasize the need for further efforts by the European commission and other institutions to establish health limits not only for Arsenic but also for other nonessential and essential elements.

Furthermore, it is important to consider the existence of organic and emerging pollutants (POPs, HAPs, PFOS, PFAS, etc.) that are becoming increasingly pervasive worldwide (Magulova & Priceputu, 2016). Given their persistence, long-range transportability, biomagnification in food chains, and bioaccumulation in humans and wildlife, their impact on individual health is becoming a growing concern (Magulova & Priceputu, 2016). The combined effects of trace elements and POPs create a complex chemical cocktail that can have synergistic or antagonistic impacts on marine life. Thus, there is a growing need for additional research to fully understand the scope of contamination which requires considering both trace elements and POPs.

### Commonly dysregulated pathways in the liver and red muscle in response to multiple-metal contamination

Liver and red muscle shared common proteome changes, such as cellular disorganization as well as functions involved in transport and stress response (Figure 5 and 7). This is in line with the positive correlation found between the functional enrichment of the two tissues in relation to mixture 1 and mixture 3 (ESM9), highlighting similar responses (upregulation). However, we noted a much more diverse response in the liver, with 17 times more pathways overexpressed compared to red muscle. Similar results have been observed in seabreams (Beauvieux et al., 2024). This discrepancy in responses between the two tissues could be related to the liver functions, which is the main organ responsible for detoxification, transformation and storage of toxic compounds in fish (Bawuro et al., 2018; Beauvieux et al., 2024).

Contaminant mixture 1 (Cu, Rb, Sr, Tl, Zn, Li, Pb and As) seemed to particularly impact cellular organization in the two tissues, as reflected by an overexpression of multiple pathways involved in cellular structure, intracellular transport and motility processes. In particular, two isoforms of cytoplasmic dynein heavy chain seemed to be affected by mixture 1 in liver (along with mixture 3). Cytoplasmic dyneins are central in intracellular trafficking, including macromolecular transport, chromosome dynamics, and the cell cycle (Paschal et al., 1987; Kardon and Vale 2009; Bawuro et al., 2018). This is in line with the significant downregulation in the pathway involved in mitotic spindle assembly that has been also detected and could indicate that trace element exposure has genotoxic effects and may repress hepatocyte division. Similarly, cytoplasmic dyneins (Table 2) were identified in bivalves as responsive to contamination (Li & Wang, 2021; Sánchez-Marín et al., 2021), which made them great candidate biomarkers of trace element contamination. Additionally, given that proper cellular organization is crucial for organismal development, the abundance of upregulated pathways associated with anatomical structure and development may suggest a response to counteract contamination-induced disorganization. Histopathological alterations in hepatocytes have already been observed, revealing the disorganization of organelles within the cytoplasm (Moreira et al., 2003) and the presence of necrosis (Javed et al., 2017) in response to metal contamination. Since hepatocytes are involved in various aspects of intermediate metabolism, unfavourable consequences to physiology may be expected as well at the cellular and organismal level (Mela et al., 2007).

Red muscle proteome response to mixture 1 displayed fewer functional diversity response than that of the liver proteome, but it reflected an alteration of the biological structure of cells and, again, the cell cycle process, as highlighted by the upregulation of the pathway “mitotic cell cycle process”. Accordingly, two of the five hubproteins identified were myosin and myomesin, which are key proteins in cell adhesion and muscle contraction. This is consistent with previous works that highlighted alterations of multiple myocytoskeleton proteins in response to contamination (Rodríguez-Ortega et al., 2003; Karim et al., 2011; Xu et al., 2019). As the contractile ability of myocytes resides in their highly organized cytoskeleton network, dysregulation here could greatly affect the locomotion of juvenile sardines, hindering their escape from predators and their foraging success.

The effects of trace element contamination on liver and muscle cytoskeleton are likely to be caused by oxidative stress (Epel et al., 2004), either through excess reactive oxygen species (ROS) generation (by redox-active metals) or reduction of antioxidant abundances and/or activities (redox-inactive metals) (Koivula & Eeva, 2010). In this study, we highlighted the upregulation of pathways such as “cellular oxidant detoxification” and “hydrogen peroxide catabolic process” in the liver and “reactive oxygen species metabolic process” in muscle. Consequently, two potential biomarkers of red muscle proteome in response to mixture 1 were identified (down-regulated gluthathione S-transferase and up-regulated hemoglobin subunit zeta, Table 3). They have both known functions in contaminant detoxification and antioxidant defences, respectively (Mollan et al., 2012). Gluthathione S-transferase was underexpressed in red muscle with increasing contamination (mixture 1), suggesting a redox-inactive potential of mixture 1 on this enzyme specifically but also highlighting that some canonical detoxification enzymes may be triggered or repressed by metal exposure, depending on their intrinsic nature (redox-active *vs* redox-inactive sensitivity).

Furthermore, trace element mixtures induced additional cellular responses aimed at safeguarding organisms from ROS-induced damage. Among these responses, a heat shock protein homologue (Dnaj homologue, Table 2) plays a crucial role by acting as cochaperone to the molecular chaperone (Hsp70). Its functions involve protein translation, promoting protein folding, translocation of polypeptides through membranes and degradation of damaged proteins (Mitra et al., 2009). In multicellular eukaryotes, programmed cell death, often referred to as apoptosis, is a prevalent stress response mechanism that comes into play when the level of stress surpasses the cell’s ability to uphold genomic and macromolecular integrity. In this study, apoptosis signalling pathway involving ADP/ATP translocase and the serine/threonine-protein kinase mTOR (Table 3 and Table 2) appeared to be affected by trace element contamination. Previous transcriptomics and proteomics studies have reported alterations in apoptotic pathways in various fish species in response to environmental contaminants (Bohne-Kjersem et al., 2009; Leaver et al., 2010; Asker et al., 2013; Yadetie et al., 2013). The serine/threonine-protein kinase mTOR was overexpressed in response to trace element contamination in the liver, which is implicated in cellular death cascades along with lipid regulation and nutrient signaling. However, besides its abundance, post-translational modifications including phosphorylation, ubiquitination, acetylation, and glycosylation appears to be key regulators of mTOR signaling and should also be considered (Yin et al., 2021). The activation of the signalling network mTOR has been associated with organic and inorganic contaminants-induced apoptosis (Chen et al., 2008; McCuaig et al., 2020). Moreover, the ADP/ATP translocase is suggested to play a significant role in the process of mitochondrial permeabilization as it can switch its function from ADP/ATP exchange through inner mitochondrial membrane to pore-formation during apoptosis (Kumarswamy & Chandna, 2009). Therefore, these two proteins could be excellent candidates biomarker for trace element effects in liver in red muscle.

### Metabolism and immune system upregulation in liver

Trace element contamination (mixtures 1 and 3) appeared to specifically affect the metabolism and transport of lipids in the liver, as reflected by the overexpression of two apolipoproteins (hubproteins). Lipids are an energy source and an essential component of cell membranes (phospholipids and cholesterol). Besides, they also play a significant role as messengers in signal transduction pathways and molecular recognition processes (Van Meer et al., 2008). Any changes in lipid metabolism would signal impairment of these pathways. It has been reported that constant energy demand leads to mobilization of triglycerides as they serve as lipid storage (Van Meer et al., 2008; Santos and Schulze 2012). The upregulation of triglyceride metabolic process and transport underlined in this study could be an evidence of their mobilization and use in repair or development of membranes damaged, for instance, under increasing oxidative stress (Van Meer et al., 2008; Santos and Schulze 2012). Their mobilization and transport may also be linked to low food intake or absorption due to gut damage or improper synthesis in the liver under trace element stress. In the absence of further research to shed light on these regulation patterns, we propose here two promising biomarkers that may indicate a dysregulation in lipid transport and metabolism in response to contamination: apolipoprotein B-100 and apolipoprotein A-IV (see Table 2) that were found upregulated in our study, but also in seahorses and atlantic cod in response to heavy metal and organic contaminant stress, respectively (Bohne-Kjersem et al., 2009; Liu et al., 2022). These proteins are important regulators of triglyceride and cholesterol metabolism (Sacks, 2006). By studying these potential biomarkers, we can gain valuable insights into the effects of metal contamination on lipid-related processes.

The sardine liver proteome also presented a clear upregulation in multiple pathways involved in immune processes. Regoli & Giuliani (2014) and Wang & Gallagher (2013) highlighted that such an upregulation could be related to trace element accumulation in the liver and lead to increasing oxidative stress. The fish immune system is known to be sensitive to environmental contamination (Si et al., 2019) and altering immune homeostasis in fish could significantly constrain their survival and development. Sustaining high levels of immune functions requires energy, which would be necessarily redirected from other mains functions, such as growth, reproduction or maintenance (energy trade-off,Moret & Schmid-Hempel, 2000).

In our experiment we did not find any correlation between body condition and contamination mixtures. This is consistent with some other studies that investigated the relationship between metal levels in the environment or fish tissues and body condition (Farag et al., 1998; Dethloff et al., 2001; Lohner et al., 2001). The measurement of body mass may not sufficiently illustrate the costs of metal exposure, since individuals can balance their resource budget and adjust their body condition over a longer period (Daan et al., 1996, Beauvieux et al., 2022). The absence of relationship could also suggest that body condition, although a great marker of nutritional stress (Brosset et al., 2015), only declines drastically when a high threshold of physiological stress is reached. Therefore, monitoring physiological processes that are more sensitive and reflect progressive changes might allow to detect early warnings of environmental challenges. Nonetheless, sardines from the Mediterranean Sea have shown a drastic decline in body condition, size and life expectancy in the past two decades (Saraux et al., 2019). The major driver proposed is the shift of sardine diet towards smaller planktonic prey observed in the Gulf of Lions due to multi-factorial environmental changes (Feuillolley et al 2020, Brosset et al., 2016), leading to lower foraging efficiency and a reallocation trade-off toward reproduction instead of survival (Beauvieux et al., 2022; Queiros et al., 2019, 2024). Consequently, individuals at the end of the reproduction period may rely on low energetic reserves to survive the end of the winter period. Adding any additional energetic or immune burden, as those highlighted in response to contaminants in this study, might emphasize their reduced chance of survival (Queiros et al., 2021).

## Acknowledgements

We are grateful to the technical and scientific crews of L’Europe for their work during PELMED survey. Furthermore, we thank Sophie Prud’homme, Roberta Bettinetti and an anonymous reviewer for their valuable inputs and comments on this manuscript.

## Data, scripts, code, and supplementary information availability

The mass spectrometry proteomics data have been deposited to the ProteomeXchange Consortium via the PRIDE (Perez-Riverol et al., 2019) partner repository with the dataset identifiers PXD037276 (liver) and PXD037313 (red muscle).

Scripts, figures 5 and 7 and other datasets are available online: https://doi.org/10.5281/zenodo.10683281

Supplementary materials are available online: https://doi.org/10.1101/2024.02.16.580673

