## Supplementary materials for "Molecular response to multiple trace element contamination of the European sardine"

**ESM1.** Quantitative proteomics of *Sardina pilchardus* liver and red muscle samples

Unless otherwise specified, all chemicals and reagents were purchased from Sigma-Aldrich (Saint-Louis, MO, USA).

**Sample preparation**

Frozen samples were ground under liquid nitrogen using a ball mill (MM400, Retsch, Eragny sur Oise, France). For liver, the resulting powder was used to extract proteins by incubation overnight at 4°C in a lysis buffer (urea 8M, thiourea 2M, Tris-HCl 1M pH 7.5, protease inhibitor cocktail 1/150). For red muscle, the incubation was performed at room temperature for 1h30 in a slightly different lysis buffer (urea 8M, thiourea 2M, DTT 2%, CHAPS 2%, protease inhibitor cocktail 1/150). After sonication for 5 minutes at room temperature, centrifugation (30s, 2000 x g, room temperature) allowed to pellet remaining cell debris. Protein concentration was determined using Pierce 660-nm Protein Assay Reagent (Thermofisher, Rockford, IL, USA). At this stage, a reference sample comprising an equal volume of all protein extracts was made, to be injected regularly during the whole experiments (liver and red muscle analyses being obviously considered as two separate experiments) and thus allow QC-related measurements.

Thirty µg of each sample were loaded onto SDS-PAGE stacking gels (4% polyacrylamide) and electrophoresis was performed for 25 minutes at 50V. After protein fixation (45% MeOH, 5% acetic acid) during 30 min, staining using colloidal Coomassie Blue (30 min) allowed visualization of a stacked protein band. The stacked protein band was excised from the gel, as well as the part of the gel above this band. Destaining was performed using 50% acetonitrile/ 50% ammonium hydrogen carbonate 25 mM, and dehydration using pure acetonitrile. Proteins were then reduced and alkylated in-gel using 10 mM DTT in 25 mM ammonium hydrogen carbonate (30 minutes at 60°C then 15 minutes at room temperature) and 55 mM iodoacetamide in 25 mM ammonium hydrogen carbonate (30 minutes at room temperature in the dark), respectively. Gel pieces were washed using 25 mM ammonium hydrogen carbonate (5 minutes at room temperature) then acetonitrile (5 minutes at room temperature), and these steps were repeated 6 times. Finally, dehydration used acetonitrile (2 x 5 minutes at room temperature), and in-gel digestion of proteins was performed overnight at 37°C using trypsin (Promega, Madison, WI, USA; 600 ng per band). After trypsin digestion, the resulting peptides were extracted at 100 rpm on an orbital shaker during 2 hours using 80% acetonitrile/0.1% formic acid in water, then they were vacuum-dried (SpeedVac, Savant, Thermoscientific, Waltham, MA, USA) and suspended in 300µL of H_2_O/acetonitrile (98/2), 0.1% formic acid for the stacked band and 30µL only for the upper band. At this stage, a set of reference peptides (iRT kit; Biognosys AG, Schlieren, Switzerland) was added to peptide extracts (1µL/9µL of liver sample; 0.7µL/6.3µL of red muscle sample) for QC-related measurements.

**Mass spectrometry analysis**

Samples were analyzed on a nanoUPLC-system (nano-Acquity, Waters, Milford, MA, USA) coupled to a quadrupole-Orbitrap hybrid mass spectrometer (Q-Exactive HF-X, Thermo Scientific, San Jose, CA, USA). The system was fully controlled by XCalibur software (v4.0.27.9; Thermo Fisher Scientific). Concentration/desalting was first performed by loading of 2 μL of sample on a Symmetry C18 trap column (100Å, 5 µm, 180 μm × 20 mm; Waters) using 99% formic acid 0.1% in water (solvent A) and 1% formic acid 0.1% in acetonitrile (solvent B) at a flow rate of 5 μl/min for 3 min. Peptide elution was then performed using a nanoEase M/Z Peptide BEH C18 column (130Å, 1.7 µm, 75 µm X 250 mm, Waters) maintained at 60 °C while applying a solvent gradient from 2% to 35% B in 88 minutes then from 35% to 40% in 5 minutes at a flow rate of 350 nL/min. To reduce carry-over, the column was washed using acetonitrile 90% for 10 minutes then regeneration of the column and a solvent blank were run in between each sample.

The Q-Exactive HF-X was operated in positive ion mode with the source temperature set to 250 °C and spray voltage to 1.8 kV. Full-scan MS spectra (375-1500 m/z) were acquired at a resolution of 120,000 at m/z 200, with a maximum injection time set to 60 ms and an AGC target value set to 3x10^6^ charges. The lock-mass option was enabled (polysiloxane, 445.12002 m/z). Up to 20 most intense peptides (at least doubly charged) per full scan were isolated using a 2 m/z window and they were fragmented using higher-energy collisional dissociation (normalized collision energy set to 27 and dynamic exclusion of already fragmented precursors set to 60 s). MS/MS spectra (200-2000 m/z) were acquired with a resolution of 15,000 at m/z 200, with a maximum injection time of 60 ms and an AGC target value set to 1x10^5^; the peptide match selection option was turned on. Peak intensities and retention times of reference peptides were monitored in a daily fashion.

**Mass spectrometry data processing**

MS raw data processing was performed in MaxQuant (v2.0.3.1) (Cox et al., 2014). Peak lists were searched using Andromeda search engine implemented in MaxQuant. The protein database contained protein sequences from a home-made annotation of the genome of *Sardina pilchardus* (TaxID: 27697; Genbank assembly accession: GCA_003604335.1; Assembly Name: UP_Spi). Briefly, coding sequences were predicted and translated using Augustus v3.4.0 software suite and protein sequences from *Alosa alosa* (TaxID: 278164; RefSeq annotation accession: GCF_017589495.1; Annotation Name: AALO_Geno_1.1) as evidence sequences. Only the longest protein per coding DNA sequence was retained. A blast search strategy using Blast+ v2.12.0 (NCBI) enabled us to attribute a name to these sequences. Blast searches first considered the species phylogenetically the closest to *Sardina pilchardus*. Using a minimum bit-score threshold set to 50, only the best blast hit found in UniprotKB (accessed in July 2022) was retained for a given *S. pilchardus* protein, allowing protein name propagation. If no satisfying blast hit was found, another blast search was performed against UniprotKB protein sequences from species phylogenetically progressively more distant to *S. pilchardus*. Blast results were validated manually. After elimination of redundancy, the database contained 58,596 protein sequences to which sequences of common contaminants were added (247 entries; contaminants.fasta included in MaxQuant), as well as decoy sequences (revert mode). The first search was performed using a precursor mass tolerance of 20 ppm, and 4.5 ppm for the main search after recalibration. Fragment ion mass tolerance was set to 20 ppm. The second peptide research option was enabled. Carbamidomethylation of cysteines was considered as a fixed modification and oxidation of methionines and acetylation of protein N-termini as variable modifications during the search. A maximum number of one missed cleavage was tolerated, and a false discovery rate (FDR) of 1% for both peptide spectrum matches (minimum length of seven amino acids) and proteins was accepted during identification. Only the proteins identified with at least two peptides were retained. Regarding quantification, data normalisation and protein abundance estimation were performed using the MaxLFQ (label-free quantification) option implemented in MaxQuant (Cox et al., 2014) using a “minimal ratio count” of one. The option “Match between runs” was enabled using a 0.7-minute time window after retention time alignment. Quantification was performed using unique peptides only. Both unmodified unique peptides and, if detected, also their modified counterpart (acetylation of protein N-termini and oxidation of methionines) were considered. All other MaxQuant parameters were set as default. The very few contaminants and reversed proteins that were identified were removed from the dataset. The mass spectrometry proteomics data have been deposited to the ProteomeXchange Consortium via the PRIDE (Perez-Riverol et al., 2019) partner

repository with the dataset identifiers PXD037276 (liver) and PXD037313 (red muscle).

**ESM2.** Mean trace elements concentration ± standard error (in mg/kg dry weight) and range measured in the 105 juveniles collected in the Gulf of Lions. For each contaminant the number of individuals with concentration above the detection limit (LOD) is indicated. Statistics are indicating only for contaminants detected in at least 50% of individuals.

|  | **N>LOD** | **Mean ± SE** | **Range (Min - Max)** |
| --- | --- | --- | --- |
| **Al** | 102 | 8.474 ± 0.920 | <LOD- 52.426 |
| **As** | 105 | 12.042 ± 0.281 | 6.397 - 18.936 |
| **Be** | 1 | - | - |
| **Bi** | 0 | - | - |
| **Cd** | 1 | - | - |
| **Cr** | 104 | 0.707 ± 0.151 | <LOD - 14.484 |
| **Cu** | 105 | 2.366 ± 0.055 | 1.449 - 5.108 |
| **Hg** | 105 | 0.046 ± 0.001 | 0.032 - 0.112 |
| **Li** | 54 | 0.371 ± 0.043 | <LOD - 2.953 |
| **Ni** | 66 | 0.379 ± 0.085 | -<LOD - 6.451 |
| **Pb** | 59 | 0.101 ± 0.008 | <LOD - 0.454 |
| **Rb** | 105 | 4.044 ± 0.055 | 2.408 - 5.328 |
| **Sb** | 19 | - | - |
| **Sr** | 105 | 3.724 ± 0.279 | 0.290 - 17.128 |
| **Ti** | 105 | 0.861 ± 0.096 | 0.167 - 7.324 |
| **Tl** | 101 | 13.060 ± 0.454 | <LOD - 20.538 |
| **Zn** | 105 | 21.374 ± 0.489 | 14.226 - 42.703 |

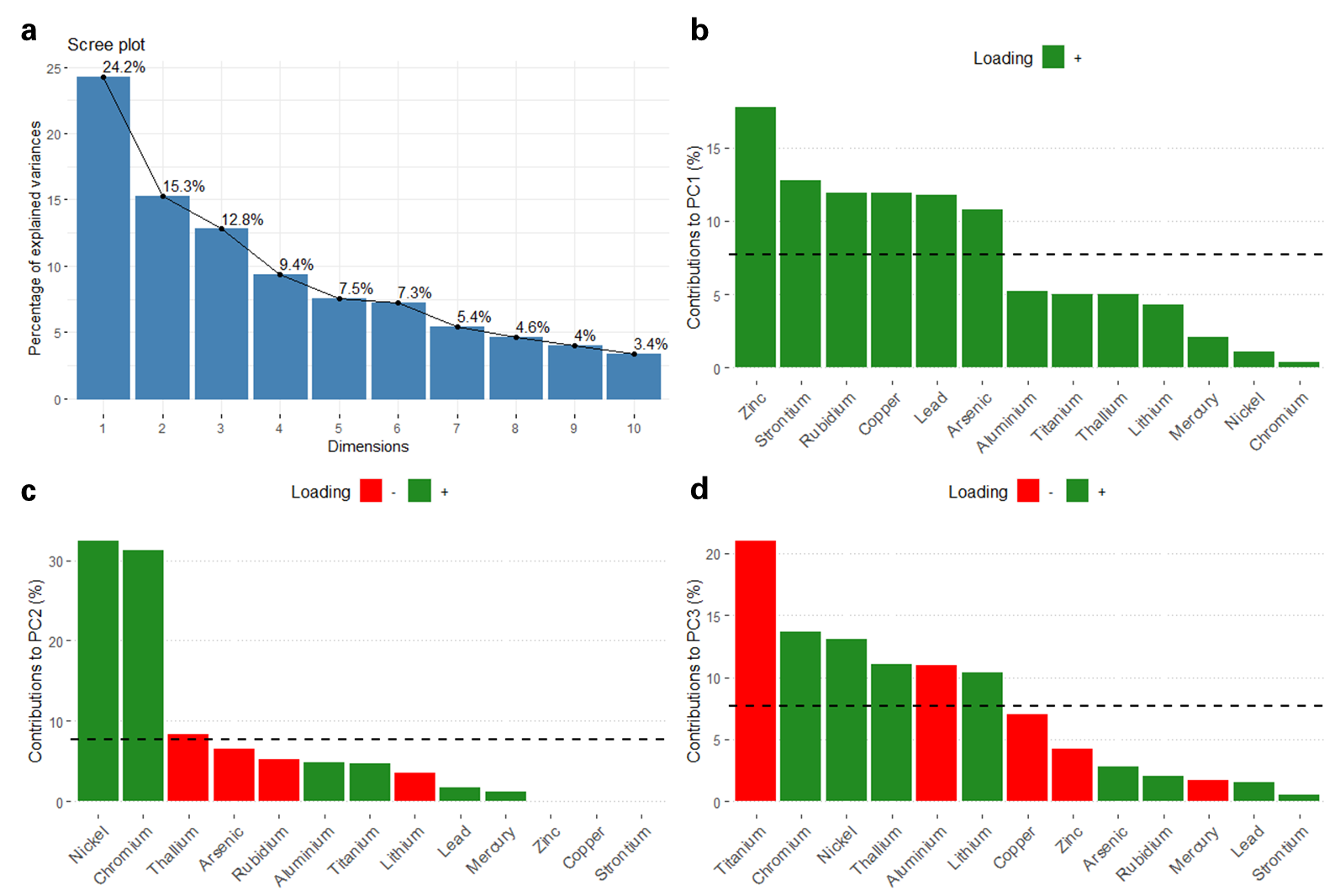

**ESM3.** Scree plot (a) and contribution of each variable to the first three principal components of the PCA on the 105 individuals. Variables loaded positively are in green while those loaded negatively are in red (b, c and d).

**ESM4.** Spearman correlation (for the 105 individuals) between distance of sampling sites from the Rhône river mouth and principal components (proxy for trace elements contamination)

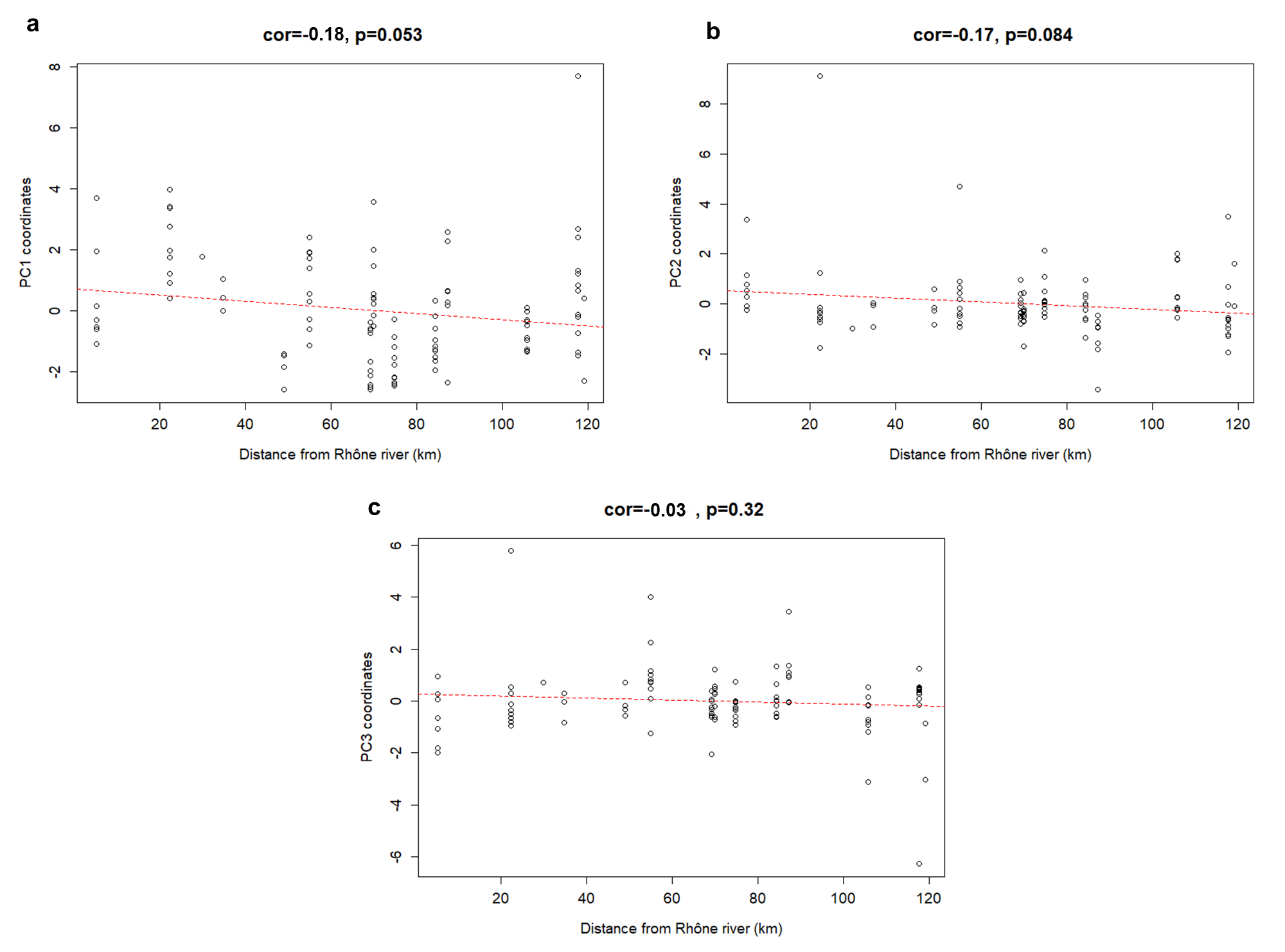

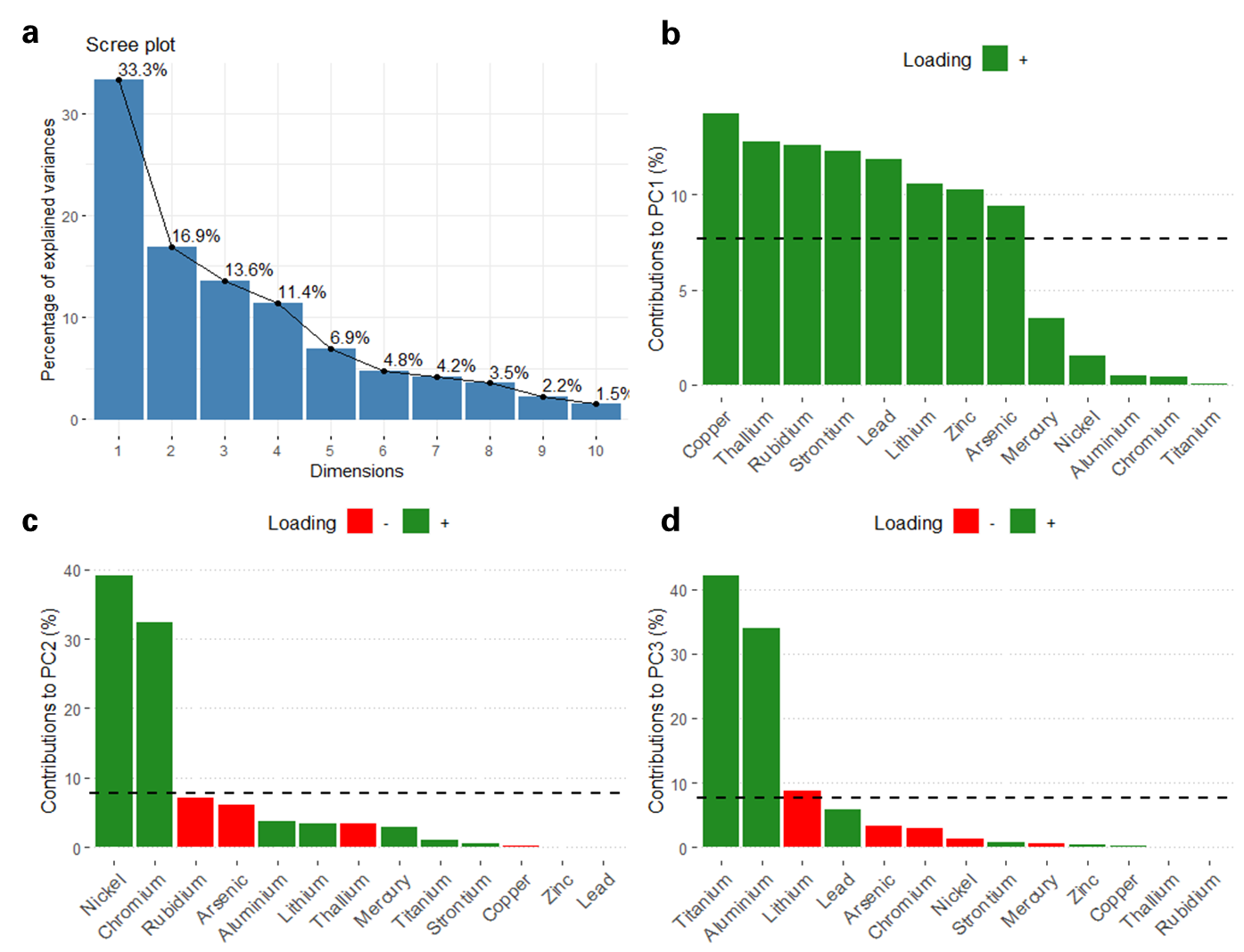

**ESM5.** Scree plot (a) and contribution of each variable to the first three principal components of the PCA on the 29 individuals selected for proteomic analysis. Variables loaded positively are in green while those loaded negatively are in red (b, c and d).

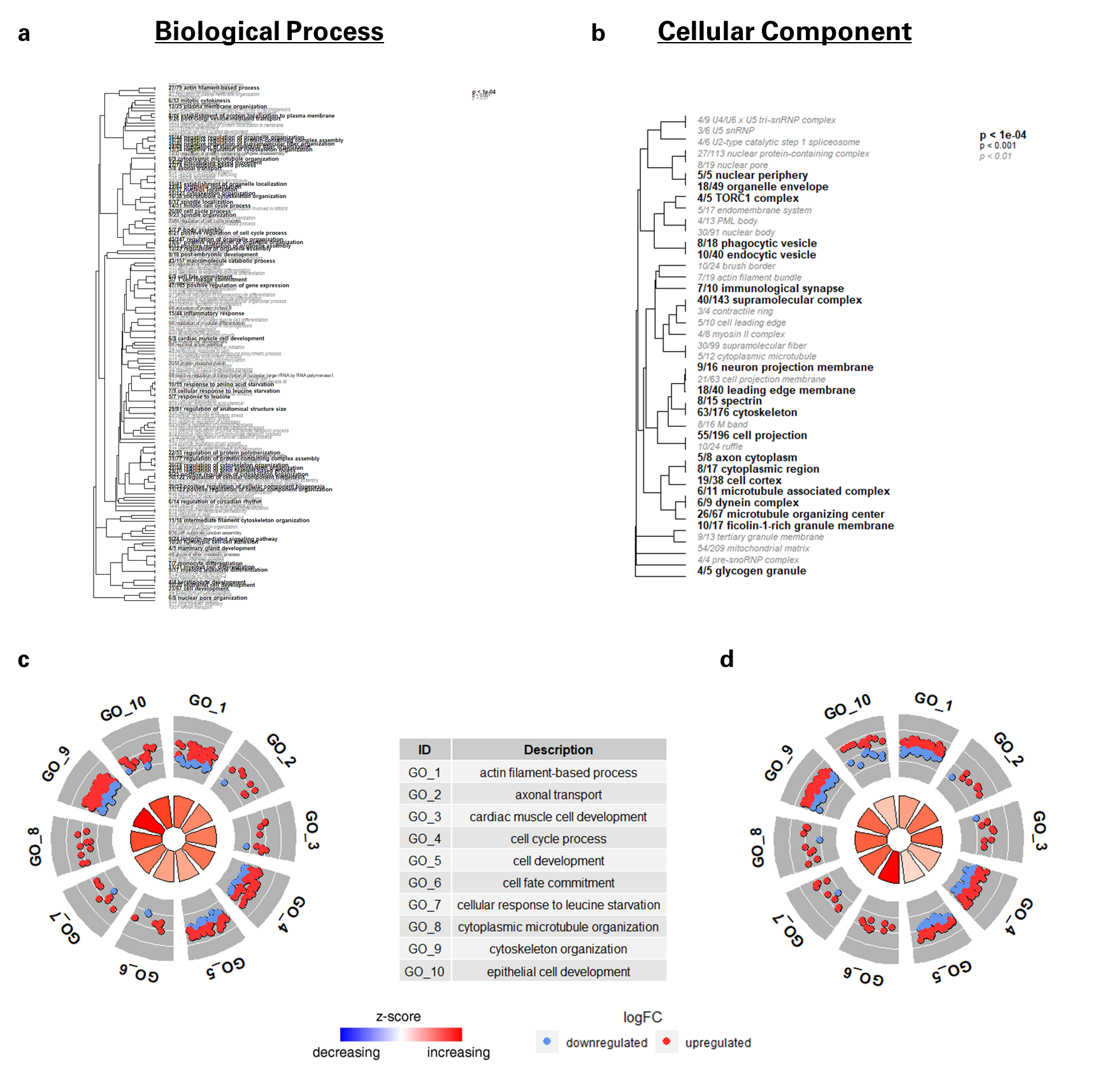

**ESM6**. Gene Ontology (GO) enrichment analysis in biological process of WGCNA protein in “turquoise” module. (a) and (b) Hierarchical clustering of enriched biological process and cellular components respectively among proteins in the turquoise module of liver proteome. Font size indicates the level of FDR-adjusted statistical significance. Term names are preceded by a fraction indicating the number of proteins within each term that are differentially regulated. GO circle plot displaying the enrichment analysis of the TOP 10 enriched biological process terms. Within each selected GO term, blue dot showed a protein underexpressed with increasing coordinates along PC1/Mixture 1 (c) and PC3/Mixture 3 (d) while red dot indicated protein overexpressed. The rectangle is coloured with the blue-red gradient according to the z-score. Z-score = (upregulated – downregulated) / √(upregulated + downregulated).

**ESM7**. Gene Ontology (GO) enrichment analysis in biological process of WGCNA protein in “yellow” module. (a) and (b) Hierarchical clustering of enriched biological process and cellular components respectively among proteins in the yellow module of liver proteome. Font size indicates the level of FDR-adjusted statistical significance. Term names are preceded by a fraction indicating the number of proteins within each term that are differentially regulated. GO circle plot displaying the enrichment analysis of the TOP 10 enriched biological process terms. Within each selected GO term, blue dot showed a protein underexpressed with increasing coordinates along PC1/Mixture 1 (c) and along PC3/Mixture 3 (d) while red dot indicated protein overexpressed. The rectangle is coloured with the blue-red gradient according to the z-score. Z-score = (upregulated – downregulated) / √(upregulated + downregulated).

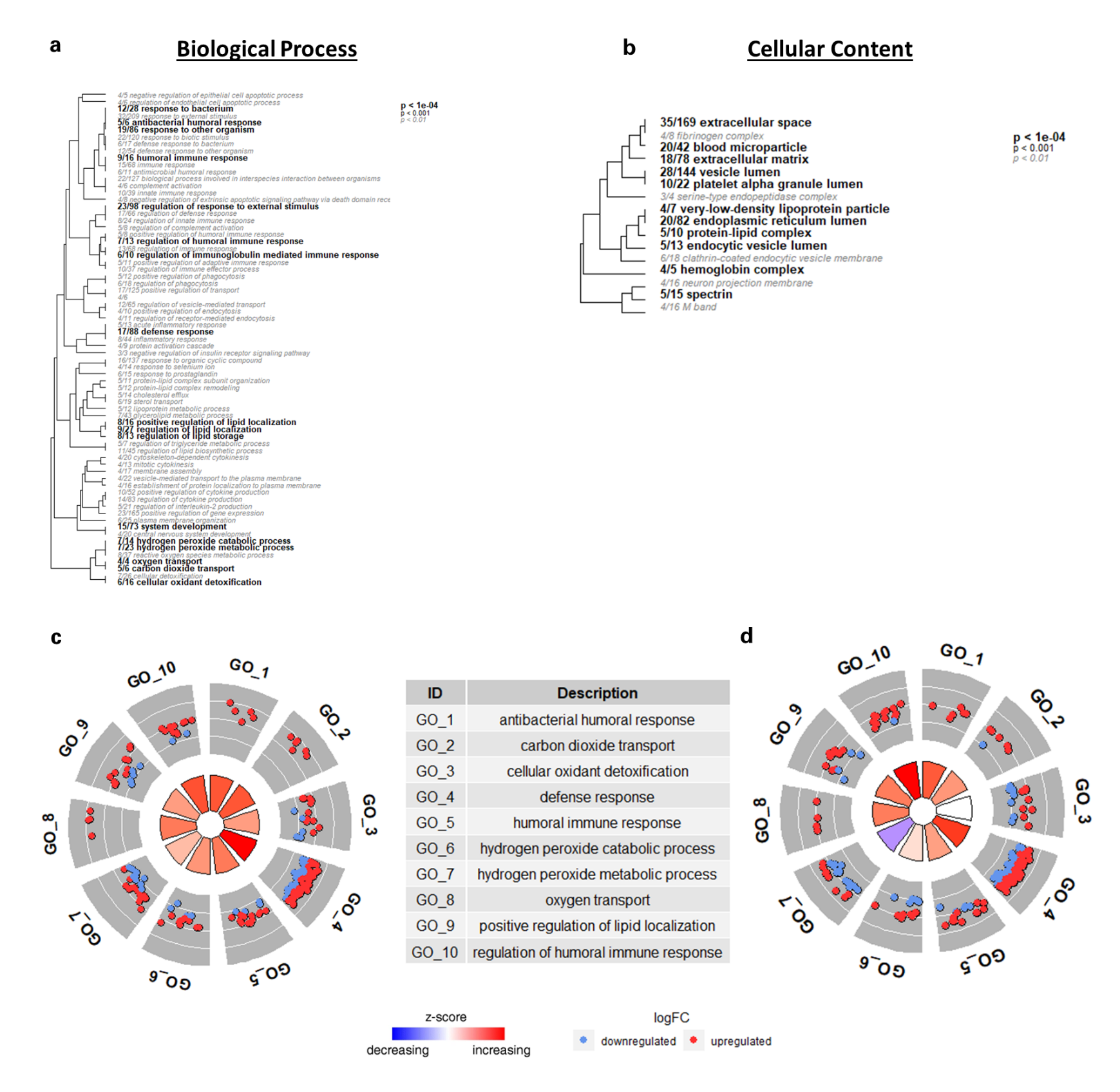

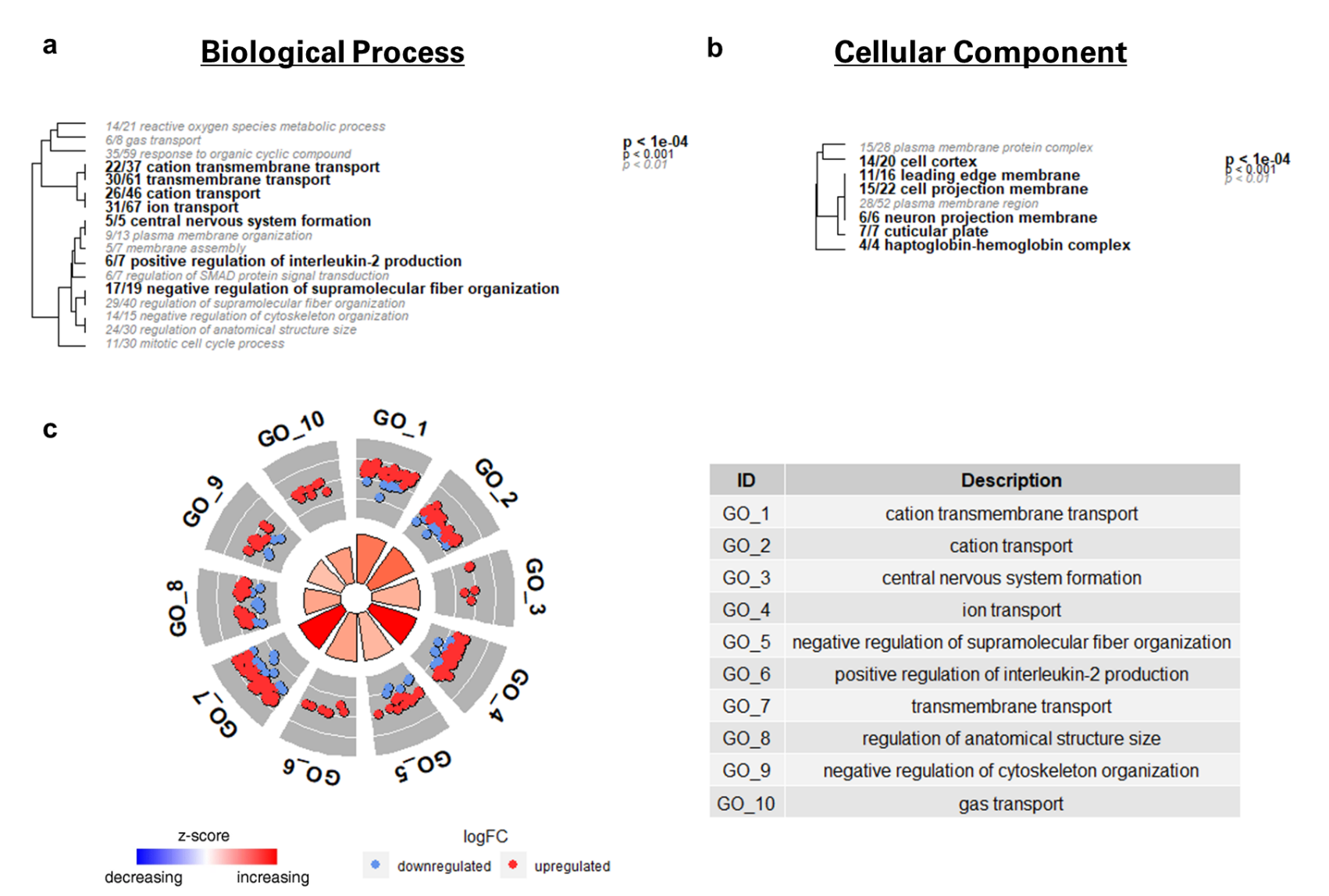

**ESM8**. Gene Ontology (GO) enrichment analysis in biological process of WGCNA protein in “blue” module. (a) and (b) Hierarchical clustering of enriched biological process and cellular components respectively among proteins in the blue module of red muscle proteome. Font size indicates the level of FDR-adjusted statistical significance. Term names are preceded by a fraction indicating the number of proteins within each term that are differentially regulated. (c) GO circle plot displaying the enrichment analysis of the TOP 10 enriched biological process terms. Within each selected GO term, blue dot showed a protein underexpressed with increasing coordinates along PC1 (Mixture 1) while red dot indicated protein overexpressed. The rectangle is coloured with the blue-red gradient according to the z-score. Z-score = (upregulated – downregulated) / √(upregulated + downregulated).

**ESM9.** Correlation of GO biological process delta ranks between liver and red muscle for Mixture 1 (a) Mixture 2 (b) and Mixture 3 (c)

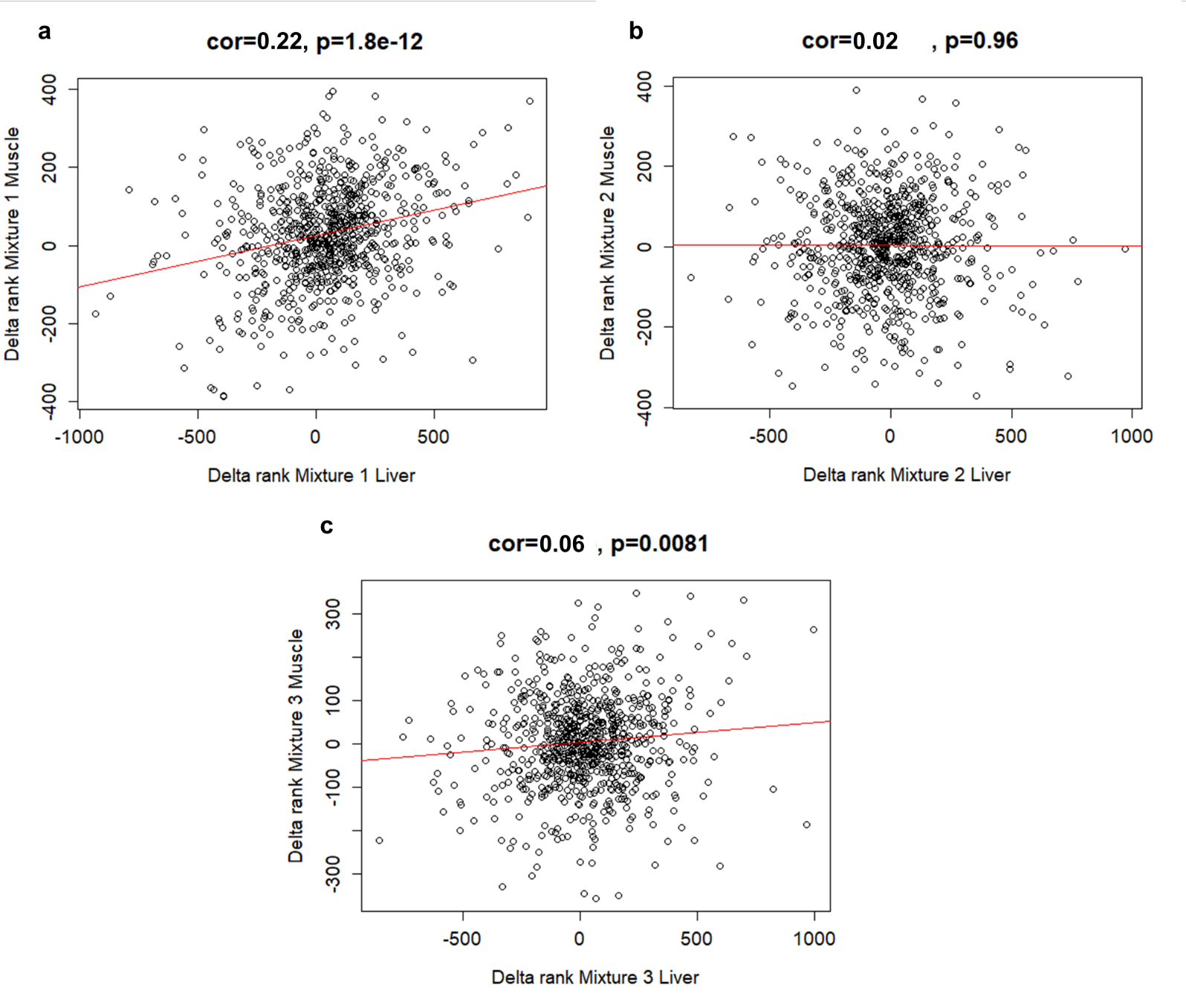

**ESM10.** Trace element concentrations in sardines (Sardina pilchardus) and close relative species muscular tissue, from the Mediterranean Sea, Atlantic and Indian Ocean. Concentrations (mean and standard deviation) are expressed as mg/kg wet weight. Some concentrations were converted in wet weight using a factor of 3.91 as suggested by Kalogeropoulos et al. (2012) (see ESM11 for further details). Number of studies in each location for each trace element are indicated below points. The dotted lines represent mean values obtained in our study.

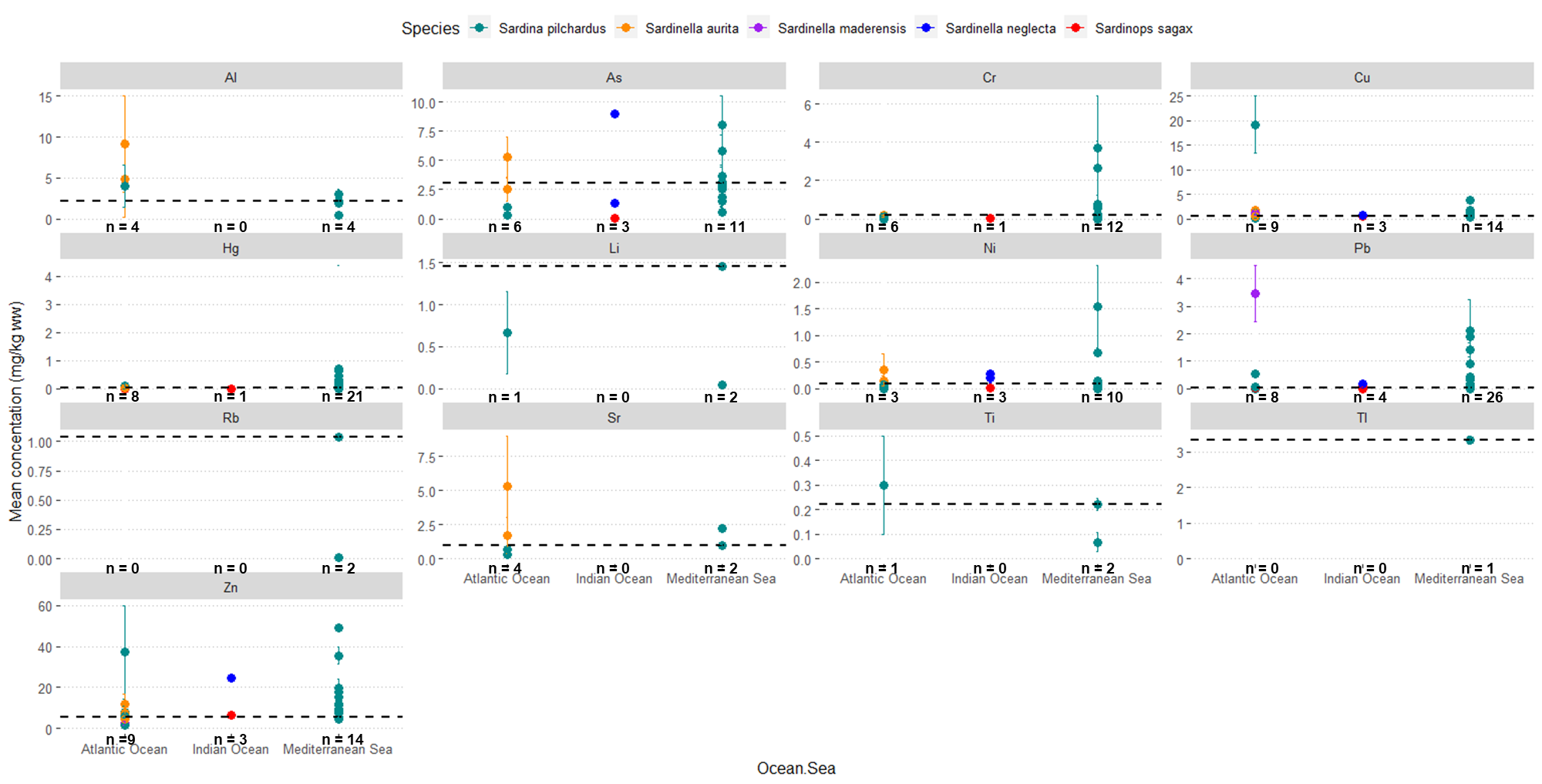

**ESM11.** Trace element concentrations in sardines (Sardina pilchardus) and other close species muscular tissue. Concentrations (mean and standard deviation/standard error, or range) are expressed as mg/kg wet weight. Concentrations indicated with (*) were converted in wet weight using a factor of 3.91 as suggested by Kalogeropoulos et al. (2012) for a moisture content of 74.4%. Last two rows represent mean and standard error as well as range of concentrations found in studies from the Mediterranean Sea only and from all studies respectively.

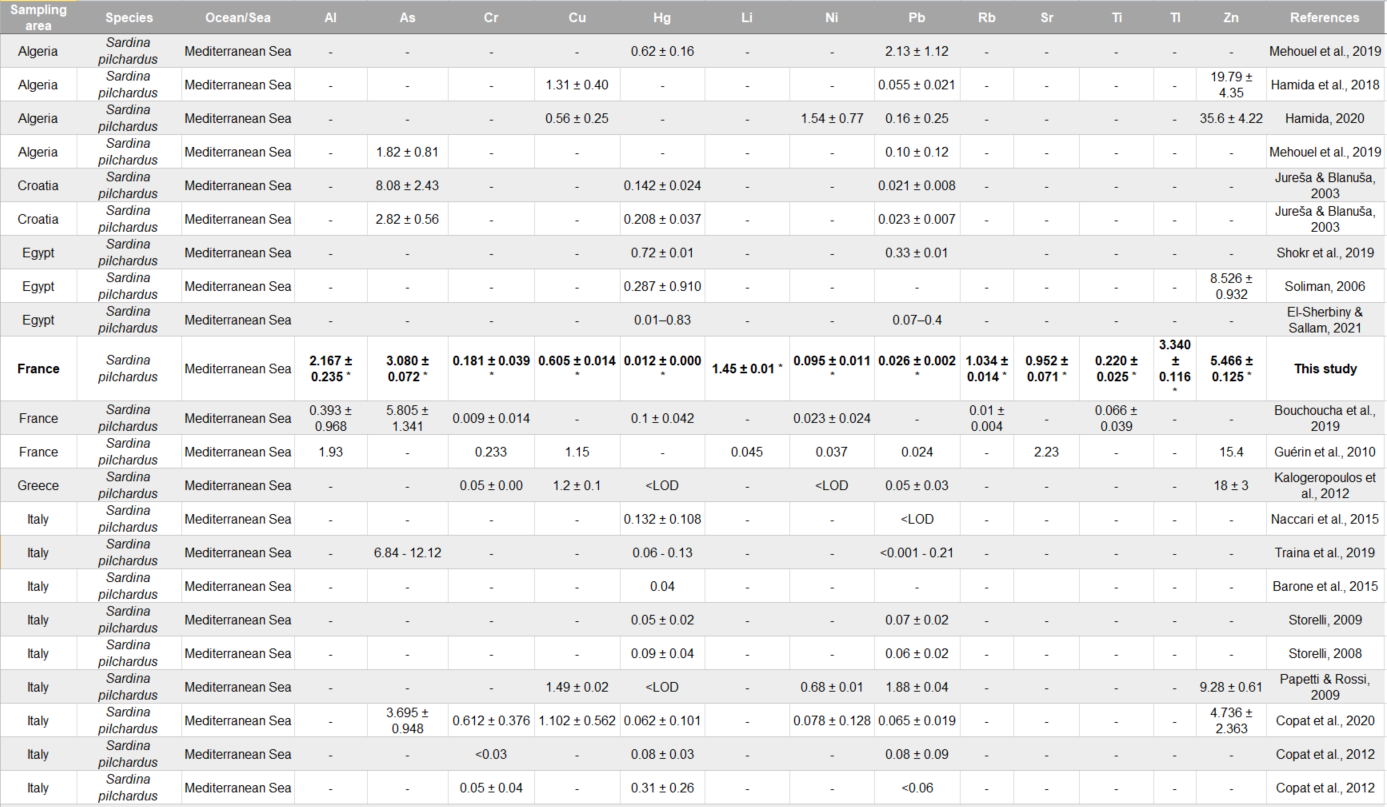

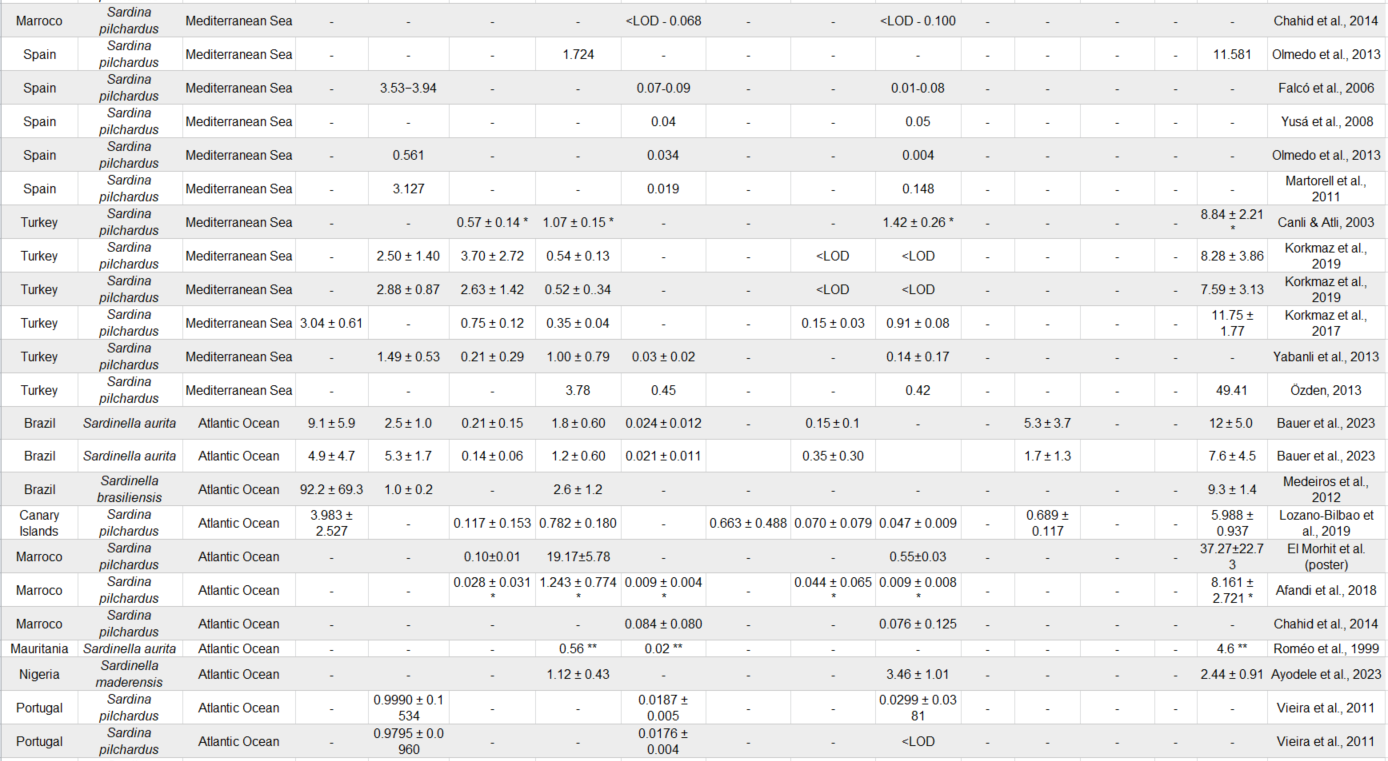

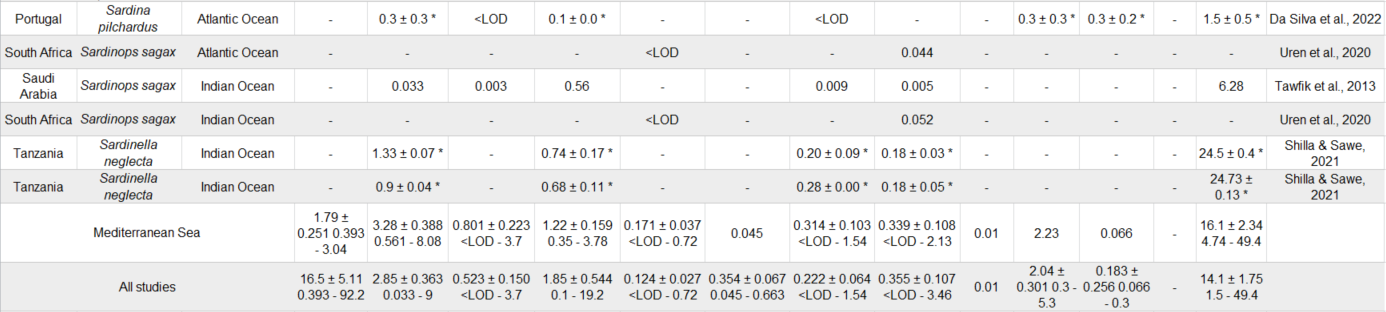
